# Time, but not reward, shapes replay-based episodic prioritization

**DOI:** 10.64898/2026.06.10.731494

**Authors:** M Tirole, É Duvelle, D Bendor

**Affiliations:** Institute of Behavioural Neuroscience, University College London; London, United Kingdom; University of Glasgow, School of Psychology and Neuroscience; Glasgow, UK

## Abstract

Why are some experiences remembered better than others? Leading theories propose that hippocampal replay prioritizes memories for consolidation according to their expected future value. We recorded hippocampal activity while rats experienced a sequence of novel environments associated with different reward values. We found that behavioral state, rather than reward, predicted replay rates during the task, and replay shifted away from reflecting current experiences as more experiences accumulated. During sleep, recent experiences were preferentially replayed, regardless of reward value. We integrate these findings into a model where each replay strengthens an episode-specific priority trace that decays over time but that can be refreshed by remote replay. We also identify a role for local replay in map stabilization, in support of the compositional mapping theory. Together, our findings demand a reinterpretation of utility-driven theories of prioritized replay and provide a framework to integrate temporal decaying episode strength into theories of memory triage and consolidation.

## Introduction

Sleep is required for memory consolidation, the process by which daily experiences are transformed into long-term episodic memories (*1–4*). However, not all experiences encountered during wakefulness are preserved. Some memories persist, whereas others fade, suggesting a process of memory triage by which the brain selectively prioritizes only a subset of episodes for consolidation (*4*). This raises a fundamental question: how are daily experiences selected for long-term storage when multiple memories compete before and during sleep?

Hippocampal replay, the reactivation of neural activity patterns representing past experiences, has been proposed as one mechanism supporting the selective strengthening of memories. During quiet wakefulness, replay is observed to occur for the current and previous behavioral contexts experienced by the subject (*5–10*). Replay during Non-Rapid Eye Movement (NREM) sleep, accompanied by coordinated cortical reactivation is hypothesized to be the mechanism that selectively consolidate these memories, with more sleep replay increasing the likelihood of consolidation, and memory triage (i.e. forgetting) as consequential to an insufficient amount of sleep replay (*5*, *11–14*). Thus, awake replay may be used before sleep to determine the relative proportion each waking experience will replay during sleep. Because replay only occurs within brief quiescent and NREM windows, is metabolically costly (*15*, *16*), and subject to homeostatic regulation (*17–19*), the economics of replay allocation for the brain is pivotal: if the brain’s “budget” for replay is constrained, then experiences must compete for access to a limited set of replay opportunities. What determines the outcome of this competition, and which daily episodes are ultimately prioritized for consolidation during sleep?

One influential answer to this replay budget allocation problem comes from value-based theories of replay (*20–22*). In prioritized memory access theory, replay is allocated according to utility, i.e. the expected improvement in future reward that would result from accessing this memory. One of the key empirical supports for this framework is the observation that replay is modulated by reward, with reward-biased replay prioritization persisting into sleep (*23–26*). As a result, reward-biased replay was interpreted as evidence that replay contributes to value learning by propagating reward information through previously experienced states, thereby supporting temporal credit assignment, value learning, and utility-maximizing planning (*9*, *21*, *27*). However, replay is also profoundly shaped by the animal’s behavioral state and recent behavioral history (*8*, *9*, *28–30*), independently of reward. This creates a significant ambiguity: the same factors that appear to prioritize replay content may also alter the states in which replay can occur. Animals may pause for longer, move less, and consequently enter behavioral states more prone to replay around high-value locations. We therefore hypothesized that previously reported reward effects may reflect increased replay *opportunity* rather than a direct influence of reward value on replay *probability*. Resolving this ambiguity is critical to understanding how memories are prioritized. In its current form, a utility-driven account of replay allocation would be challenged, with broad implications for theories linking replay to value learning and reward-based planning.

Moreover, memory triage cannot be fully understood by studying isolated episodes, where competition is minimal. Instead, it requires examining how the replay is allocated across multiple, *contextually distinct*, experiences within a single wake-sleep cycle. Although replay is often implicitly treated as a limited resource (*22*, *31*, *32*), direct evidence for the resulting trade-off between competing episodes remains elusive. Furthermore, most studies have focused on familiar tasks in single environments, where replay can be related to subsequent recall or behavioral performance. While this approach has been instrumental in establishing links between replay and memory, it leaves largely unexplored a central feature of competition across episodes: time. Episodes within a wake bout necessarily differ in their temporal proximity to sleep, a factor known to strongly influence memory retention (*33*, *34*). Any theory of replay allocation must therefore explain not only how replay is distributed across competing episodes, but also how temporal position shapes the outcome of that competition.

If previous episodes compete with the current experience for a limited replay budget, then replay allocation may do more than just select which episodes are later consolidated. Indeed, replay has also been proposed to help stabilize hippocampal representations (*35–37*), with recent compositional accounts of replay proposing that new representations can be constructed by combining episode-specific features with reusable structural elements from prior experience (*38*). Allocating replay toward or away from the current experience could influence whether a stable, episode-specific, representation is formed at all. Consequently, replay of previous episodes may indirectly shape the construction of new neural representations (*39*).

In the present study, we tested how replay priority of competing memories is allocated by the hippocampus, accounting for both value and history. We investigated three fundamental aspects of how replay may support memory triage: (i) does reward value directly bias replay selection, as proposed by utility-maximizing theories of replay allocation, or does it expand replay *opportunity* by altering behavioral state, challenging this view?; (ii) whether replay allocated under competition between episodes is shaped by recency - a largely unexplored dimension of memory triage; (iii) does replay’s role in memory triage go beyond episode selection, or does it also contribute to stabilization of hippocampal representations during experience, linking replay allocation to both episode consolidation and formation?

## Results

We recorded hippocampal activity from five food-restricted male rats as they ran back and forth on three novel two-meter tracks presented sequentially each day, with rest periods before, between, and after the track runs (Fig.1A). Each track provided either a HIGH value reward (chocolate milk) or a LOW value reward - created by diluting the same liquid 1:1 with water. Rats shuttled between the two track ends for 15 minutes to collect 0.1 mL of liquid at each reward site. Between track runs (“episodes”), they rested in a remote enclosure (“sleep pot”) for 10 minutes. At the beginning and end of each session, animals rested for up to 60 minutes and 90-120 minutes, respectively. Novelty was maintained by altering track geometry and local and global cues each day, with all three tracks sharing the same geometry within a given session.

**Fig. 1:**
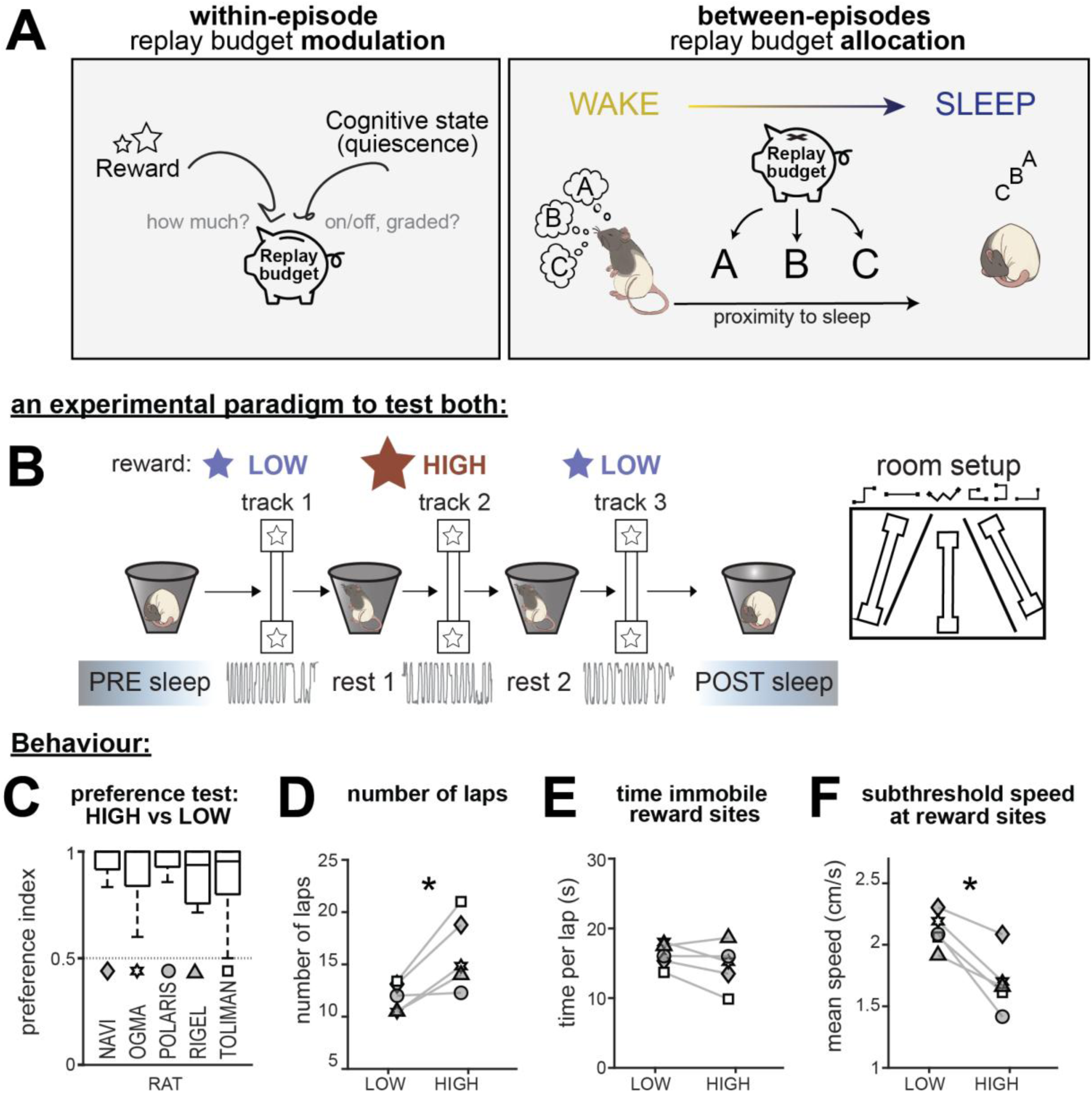
replay budget accumulation and allocation: experimental design and behavior. **A**: schematic of the working framework: the replay budget for an episode is set during its experience. Here, it can be increased by rewards value and/or the animal’s behavioral state. The latter may act as an on/off switch or display a more graded influence. The replay budget is allocated during wake between competing episodes. This allocation, coupled with proximity to sleep may influence subsequent consolidation. **B**: Experimental design. Each session, three separate tracks are experienced. Rats run back and forth for 15 minutes to collect reward on the end platforms. Two tracks are associated with one reward value (HIGH or LOW) while the third has the other reward value (LOW or HIGH), counterbalancing reward and recency. Rats are left to rest in a separate sleep box at the beginning (PRE) and end of the session (POST), and in between each track experience (Rest1,2,3). **C-F**: behavioral measures demonstrate reward sensitivity. Rat averages for LOW vs HIGH tracks. Rats used for replay analyses have filled in symbols. Denoted by (*) significant contrast HIGH-LOW obtained from GLMMs ∼ reward × track + (1 | rat/session). See corresponding Table S1 and supplementary text. **C**: Reward preference assay. All rats preferred the HIGH reward (pure chocolate milk) over the LOW reward (1:1 dilution with water), indifference point is 0.5. **D**: number of laps ran on the track. Main effect of Reward (*F*(1,60.22) = 22.479, *p* < 0.001). **E**: time spent immobile at reward sites, per lap (main effect of Reward *F*(1,1203.69) = 3.048, *p* = 0.08). **F**: degree of immobility at reward sites as measured by the mean speed for periods when speed < 5cm.s^-1^ (main effect of reward *F*(1,1218.67) = 89.62, *p* < 0.001).

### Behavioral state, not reward value, expands within-episode replay opportunities

#### Rats demonstrate sensitivity to reward value

Before recordings, rats underwent a reward preference assay with unrestricted access to both reward types. All animals consistently preferred the HIGH reward, consuming very little of the LOW reward (Fig.2A).

**Fig. 2:**
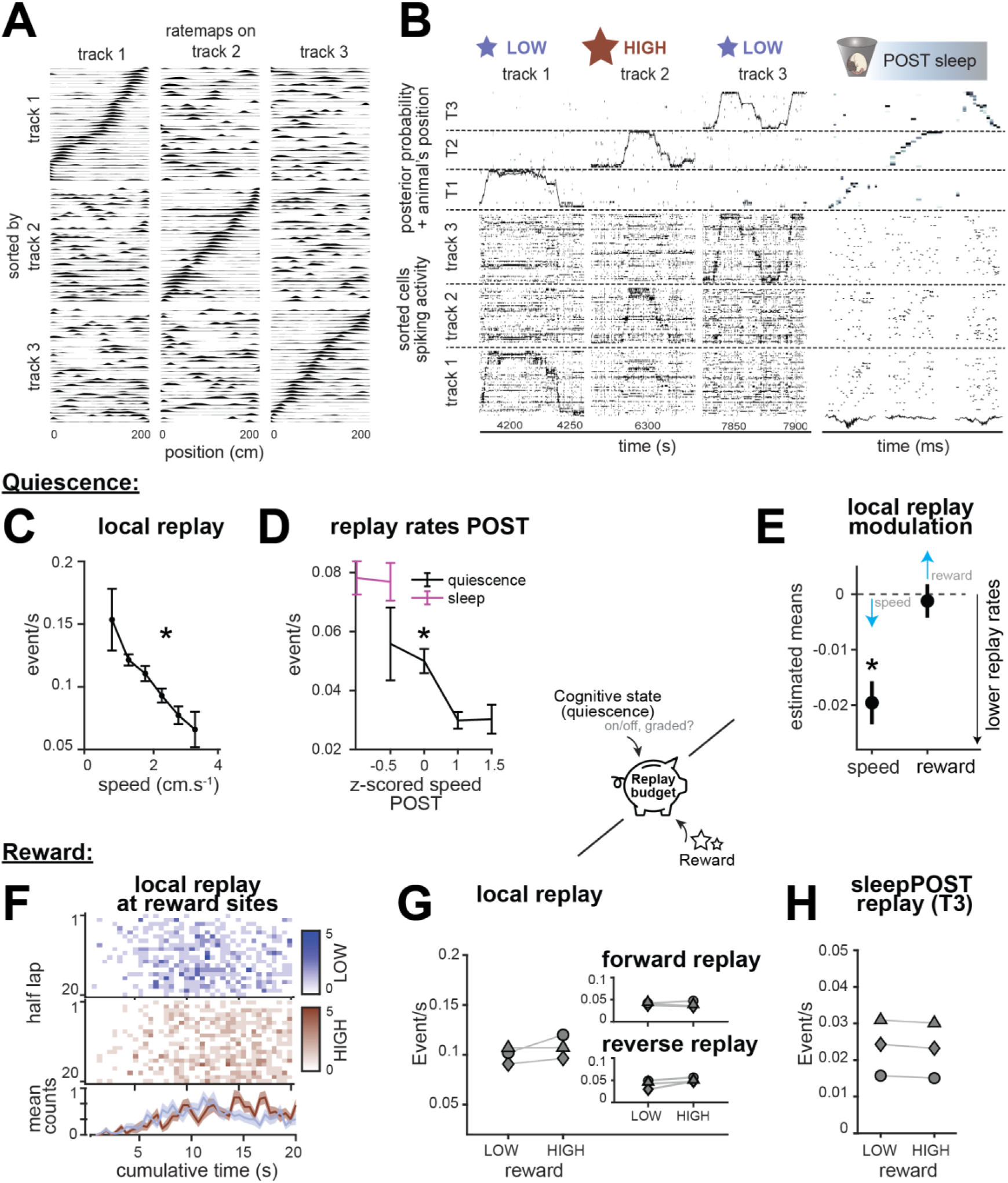
Within-episode replay allocation: behavioral state, not reward value, increase within-episode replay rates. A-B: Example hippocampal neural activity and decoding. **A**: Normalized rate maps (for visualization only) of all spatially tuned cells, calculated for each episode (rows), and sorted by their peak firing rate on another episode (columns). Note the significant amount of remapping between episodes, with on-diagonal pattern only preserved for same-same comparisons. **B**: Raster plots of spiking activity, repeated for Track 1,2 and 3, sorted by the preferred firing location of each cell on the corresponding episode. On top, decoded position from neural activity with the true location of the animal superimposed. Darker colors indicate a higher posterior probability. Right: three examples of decoded replay events during POST. Bottom: LFP traces at the time of replays. **C-E:** increased replay rates track the degree of quiescence. **C**: mean local replay rates as a function of the mean subthreshold speed (v < 5 cm.s^-1^) per lap, averaged across sessions (*β* = −0.034, *t* = −11.724, *p* < 0.001 ). The lower the speed, the higher the replay rate, see Table S8. **D**: mean replay rates during POST as a function of the mean z-scored speed, split between quiescent (in grey) and sleep states (in magenta), averaged across sessions (*β*_*quiet*_ = −0.02, *t* = −3.89, *p* < 0.001). Replay rates increase with lower speeds in the sleep pot. See Table S8. **E**: estimated mean and standard error for the main effect of speed and reward on local replay rates, derived from the GLMM rate per lap ∼ speed (v < 5 cm.s^-1^) + lap number + reward × track + (1 | rat/session). Higher speeds are linked with lower replay rates (*β*_*speed*_ < 0, main effect of speed F(1,583.952)=26.043, p<0.001), while reward does not influence replay rates (*β*_*reward*_∼ 0, no main effect of reward *F*(1,700.747) = 0.151, *p* = 0.698), see supplementary Table S4. **F-H:** replay rates are not modulated by reward value. **F**: sum of local replay counts over all rats and sessions, binned per lap and cumulative time spent immobile (v < 5 cm.s^-1^) at reward sites, split for LOW and HIGH value tracks; 0 indicates the first data point spent below the speed threshold at the reward zone. **G-H**: mean replay rates for LOW and HIGH value tracks, per rat. **G**: local replay rate (no main effect of reward *F*(1,700.747) = 0.151, *p* = 0.698), inset top: local forward replay (no main effect of reward *F*(1,693.059) = 2.329, *p* = 0.127), inset bottom: local reverse replay (no main effect of reward *F*(1,678.729) = 1.075, *p* = 0.3). **H**: offline replay of track3 during sleep POST (no main effect of reward *F*(1,46.001) = 0.18, *p* = 0.674). All replay rates are computed per second of time spent at low speed and/or asleep.

During recording sessions, rats clearly distinguished reward values: they ran faster on HIGH compared to LOW episodes (Fig.S1F, Table S1), completed more laps for HIGH than LOW in the same timeframe (Fig.2B, Fig.S1A, Fig.S1F, Table S1), and showed a small but robust decrease in subthreshold speed at the HIGH reward sites (Fig.2D, Fig.S1G, Table S1).

Critically, time spent in replay-permissive quiescent states was largely matched across conditions, in contrast to prior experimental designs (Fig.2C, FigS1B, Table S1). While quiescence depth was slightly stronger in HIGH episodes due to lower subthreshold speeds, our experimental design minimized gross differences in replay opportunity arising from unequal stopping durations. Similar results were obtained using a conservative immobility threshold (v < 2 cm·s⁻¹, Fig.S2, Table S2). Full statistical interpretation of results is available in supplementary.

This places us in a strong position to test whether reward value alters replay probability itself, as predicted by utility-maximizing theories of replay allocation.

#### The degree of quiescence correlates with replay rates beyond a simple switch ON/OFF mechanism

Out of five rats, we recorded a sufficiently large populations of CA1 place cells in three animals to perform replay detection and decoding. Briefly, a naïve Bayes decoder was used to infer the animal’s position from hippocampal activity. Decoded sequences occurring during low-speed bursts of activity were compared against three types of sequence-based shuffle methods to identify significant replay events, which were then assigned to a single episode (i.e. track). Forward and reverse replay were detected using directional rate maps (see Methods, Fig.2A-B).

The following terms are used throughout: we refer to replay of the current episode (track) while the rat is physically on that specific track as *local replay;* replay of a different episode while physically on the track as *online remote replay*, and finally, any replay while physically in the sleep pot as *offline replay*; the latter is further divided into awake and sleep replay.

Our behavioral analyses established that replay-permissive states were broadly matched across reward conditions. Although replay is generally associated with quiescence, hippocampal state transitions are not strictly binary (*40*). We therefore asked whether replay opportunity itself varies continuously with the degree of quiescence by quantifying the relationship between replay rate and subthreshold speed.

Local replay rates decreased monotonically with increasing subthreshold speeds (Fig.2C, Table S4), a relationship that also held for offline replay rates during POST (Fig.2D, Table S4). Although these initial linear fits revealed a continuous relationship between quiescence and replay rate, they do not account for experience-dependent changes in replay rates across laps ((*32*, *41*), Fig.3A).

**Fig. 3:**
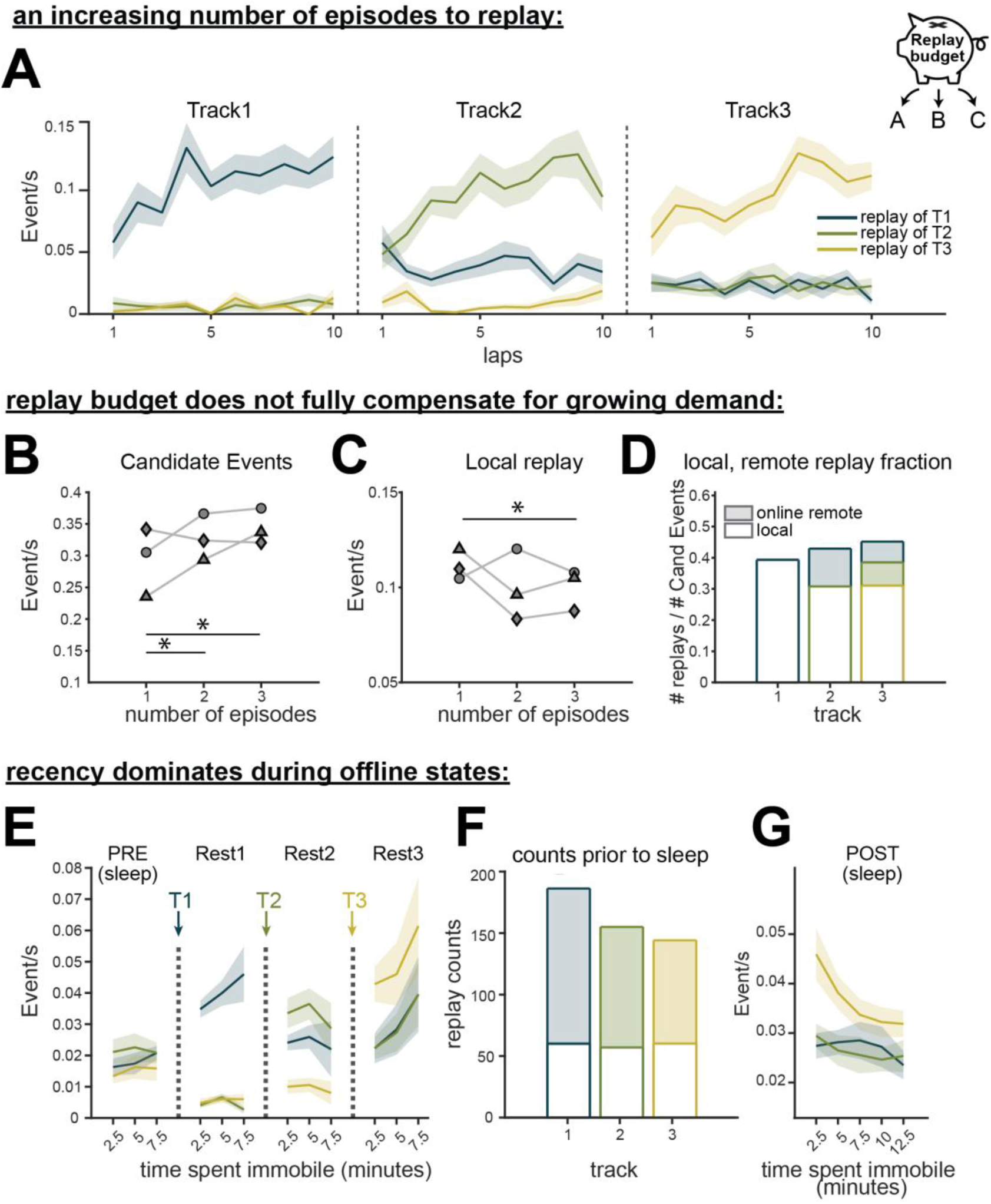
While competition between multiples experiences limits local replay, Recency dominates online and offline replay. **A**: Replay rate as a function of the number of laps during an episode, note how local replay is always highest and increases throughout the session, while replay of past episodes (remote) appears once they have been experienced. **B-C**: event rates as a function of the number of episodes in a session. (*) denotes significant contrasts TrackY-TrackX obtained from GLMMs ∼ reward × track + (1 | rat/session). Lines indicate which contrasts are significant, e.g. Track3-Track1 in C. **B**: Candidate replay events (main effect of track *F*(2,693.61) = 10.30, *p* < 0.001). **C**: local replay events (main effect of track *F*(2,691.59) = 3.792, *p* = 0.023 ). **D**: fraction of replay events to the number of candidate replay events as a function of the number of episodes, color coded by content, shaded portions represent fractions of remote events. **E,G**: offline replay rates evolution over minutes of immobility (or when indicated, sleep) in the sleep pot, color coded by content for sleep PRE, Rest1, Rest2, Rest3 and in G sleep POST. Replay of the most recent episode is most prevalent during Rest1-3 and sleep POST, but at chance during sleep PRE. In G, replay rates of all episodes gradually decreased with time spent asleep (*β* = −2.27 × 10^−5^, *t* = −4.813, *p* < 0.001). See Table S8 **F**: mean replay counts for episodes 1, 2 and 3 prior to sleep per session, shaded portion indicates fraction of remote events.

We therefore fitted a generalized linear mixed-effects model (GLMM) with local replay rate as the outcome, subthreshold speed and lap number as continuous fixed effects, and session nested within rat as a random intercept (see Methods). In this model, local replay rates increased with lap number but decreased with subthreshold speed (Fig.2E, Table S4).

These results indicate that replay is not simply switched on once a quiescence threshold is crossed but varies monotonically with behavioral state. This result is relevant to interpreting replay rate differences across conditions, as changes in quiescence could confound apparent effects of reward.

#### Reward value does not increase the replay rates of the current episode

If we account for replay opportunity varying continuously with behavioral state, do we observe reward value directly biasing replay selection, as proposed by utility-maximizing theories of replay allocation? If so, HIGH-value episodes should still be replayed more even after accounting for replay-permissive behavioral states.

Previous studies reported elevated replay rates during HIGH-reward immobility bouts (*23–26*, *42*). We first reproduced the visualization from Ambrose et al., plotting replay counts as a function of cumulative time spent at reward sites. In contrast to these previous reports, replay rates in our dataset appeared similar for HIGH and LOW reward episodes, where time spent at reward sites was broadly matched across conditions (Fig.2F).

To quantify these effects, we used a GLMM that extended our preceding quiescence analysis. Reward value and track identity were added as categorical fixed effects, while retaining subthreshold speed, lap number, and the same random-effects structure. This approach allowed us to explicitly partition out the contribution of quiescence and novelty before testing whether reward explained any remaining variance in replay rates. Importantly, reward value did not significantly modulate online replay rates, whether replay rates were measured across the full track or restricted to reward sites, and regardless of whether we considered candidate events, local replay, forward replay, or reverse replay (Fig.2G, Fig.S4, Fig.S5, Table S4, Table S5, supplementary text).

We additionally tested whether reward value modulated offline replay during rest and sleep. Consistent with our results for online replay, offline replay rates during inter-episode REST, POST, and sleep POST epochs did not differ between reward conditions (Fig.2H; Fig.S4; Table S4). Anecdotally, a small, non-significant elevation in replay rate during Rest3 for Track 3 HIGH relative to Track 3 LOW was linked to a greater immobility and shorter latency to sleep onset following Track 3 HIGH sessions (Fig.S1H,I), again, supporting the idea that replay allocation remains more closely linked to behavioral state than reward value itself.

Finally, neither current reward value, previous reward value, nor their interaction significantly influenced local replay rates, on track 2 or track 3 (see supplementary text, Table S9), ruling against the possibility that replay rates were modulated by reward history and by relative comparisons between rewards within a session.

#### Reward value does not modulate replay quality

Although replay *rates* were unaffected by reward value, reward value could still influence other features of replay. We therefore tested whether HIGH-reward episodes were replayed with greater fidelity (*43*), which could lead to better subsequent consolidation. We compared the weighted correlation score of local (bidirectional), forward, and reverse replay events for LOW and HIGH value replay content. We found no significant effects of reward, recency, or their interaction for any replay type (Fig.S6, Table S6). These results indicate that reward did not alter the fidelity of replay sequences.

Together, these findings argue against a strictly utility-maximizing account of replay prioritization and instead suggest that behavioral state primarily determines the amount of replay of the current episode.

#### Pressure from multiple experiences creates competition

If awake replay prioritizes memories for later consolidation, then replay of the current episode should increasingly compete with replay of prior episodes as additional experiences accumulate across the wake bout.

We observed that during each episode, local replay dominated (Fig.3A). Remote replay of previous episodes could also be detected throughout each experience, accounting for about a third of online replay events (Fig.3D). Consistent with prior work, local replay rates increased over the first few laps of each episode (Table S4), potentially reflecting progressive stabilization of the current hippocampal representation.

As additional episodes were experienced, the demand placed on replay increased (Fig.3B) without a corresponding increase in quiescence (Fig. S1). Consistent with replay being rate limited, while candidate replay event rates increased modestly with episode number (Fig.3B, Table S4), they failed to offset the growing prevalence of remote replay. Consequently, local replay rates declined across episodes and were significantly lower on Track 3 than on Track 1, with a similar trend for Track 2 (Fig.3C, Table S4). Local replay therefore occupied a progressively smaller fraction of replay events, while replay of previous episodes consumed an increasing share of the replay budget (Fig.3D).

#### In the absence of other factors, recency dominates replay allocation during offline states

Unlike online replay, offline replay is not directly anchored to ongoing sensory cues that may bias replay towards the current environment (*6*, *7*, *9*, *44*). Despite the progressive reduction in local replay rates across the sequence of episodes, replay during subsequent offline states (REST) was dominated by the most recent experience (Fig.3E, Table S4). Moreover, the relative replay rates of different episodes established online appeared to be preserved offline, an effect we return to in a subsequent section.

We next asked whether this recency bias persisted into sleep. One possibility is that sleep prioritizes the episode with the strongest or most stable representation. In our dataset, this would predict preferential replay of Track 1, which accumulated significantly more replay before sleep than later episodes (Fig. 3F). Instead, consistent with the REST periods, the most recent episode dominated sleep POST, and replay rates decreased with time spent asleep (Fig. 3G). Therefore, in our dataset, temporal proximity to sleep outweighed cumulative prior replay in determining replay allocation during offline states.

These findings identify recency of experience as a major determinant of replay-based memory triage. In the absence of strong competing factors, replay allocation persists from the currently experienced episode during online behavior as the most recent episode during offline states and sleep.

### Local replay is associated with the stabilization of emerging compositional maps

The preceding analyses showed that replay allocation shifts dynamically across current and previous episodes during wakefulness and sleep. This raises a further question central to memory triage: what are the consequences of allocating replay locally to the current episode? Recent models propose that hippocampal representations are constructed compositionally, by binding reusable relational building blocks into the current environment configuration (*38*). In line with this perspective, place field stabilization should correlate with the dynamics of local replay, possibly for integrating newly generated rate maps within a pre-existing, schema-like structure shared between episodes.

To test this idea, we classified cells according to their participation in past and current episode representations. Cells active only in previous representations were defined as *past cells*, cells active in both previous and current representations as *shared cells*, and cells active in the current representation but not in previous representations as *newly recruited cells* (Fig.4A). To avoid any ambiguity in cell classification for Track 1, related data were excluded from analyses in panels Fig.4D-J.

**Fig. 4:**
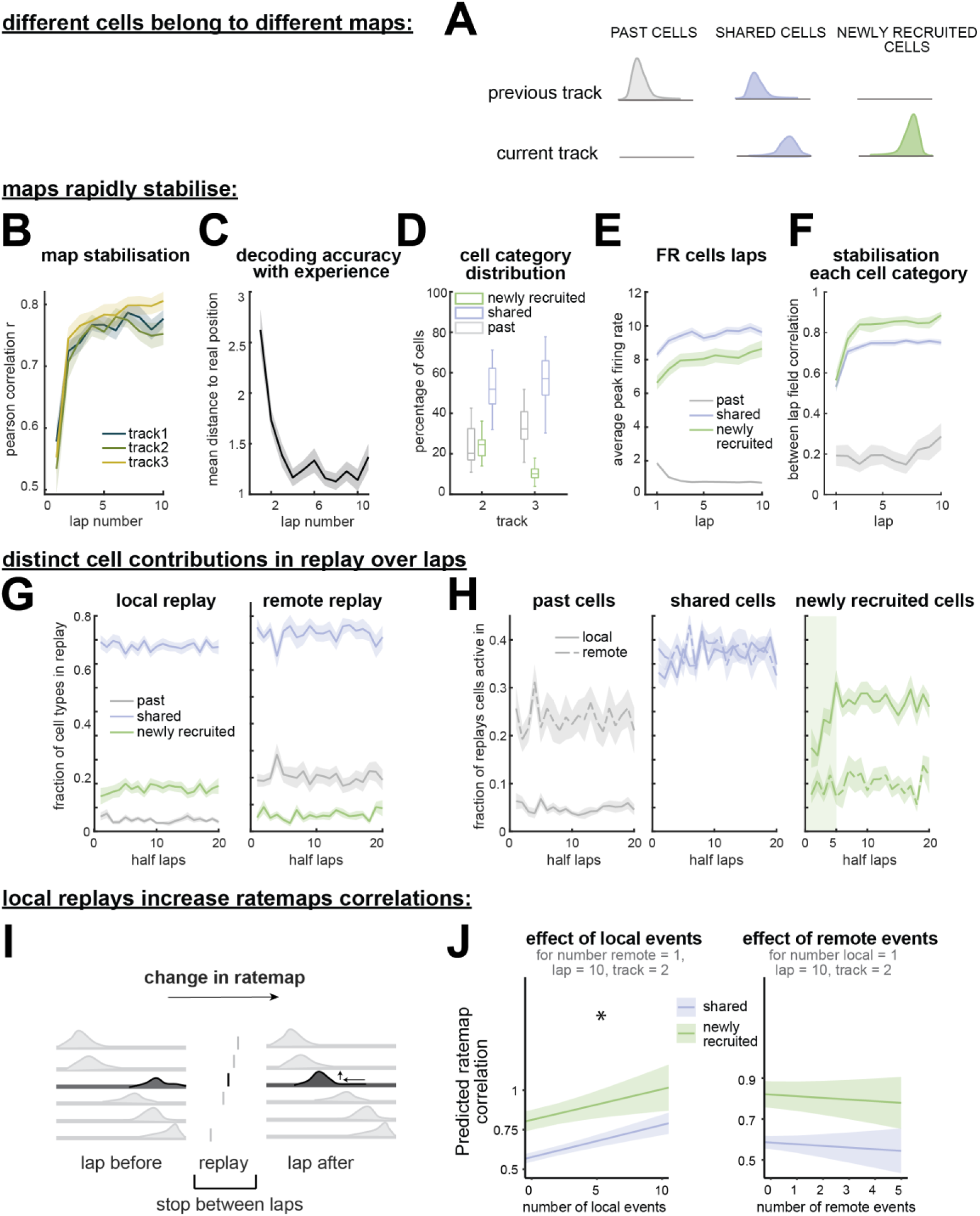
Compositional replay is linked to the stabilization of spatial maps. **A**: Categorizing cells active during an episode. **B**: Pearson correlation (r) of rate map population vectors between consecutive laps of a novel environment plateaus rapidly, independently of episode number. **C**: distance between the animal’s location and the maximum amplitude location of the posterior, as a function of the number of laps during an episode. **D**: distribution of the number of cells in each category for each episode. **E**: peak in field firing rate per lap, for each cell category (see schematic in A). **F**: Pearson correlation (r) of rate maps between laps, for novel and active shared cells. Novel cells stabilize to a higher degree than shared cells, while past cells never stabilize, as expected since they were defined to not integrate the new representation (interaction cell category x lap number *F*(1,9326.7) = 8.29, *p* = 0.004). **G**: fraction of cells represented by each category during local and remote replay events, as a function of the number of laps during an episode. **H**: fraction of local and remote replay events as a function of the number of laps in an episode, split by cell category. (left: past cells, middle: shared cells, right: novel cells). Unlike the other two cell categories, novel cells exhibit a gradual increase in participation in local replay events over the first 5 laps. **I**: schematic of how rate maps change before and after replay events for cells active during said events. **J**: predicted changes in Pearson correlation (fisher transformed) from the model: rate map correlation (zF) ∼ track + cell category × lap number + cell category × number of replay local where cell is active + number of remote replay where cell is active + (track + lap number + cell category || cell id). Once controlling for the effect of experience on the track (lap number) and track identity, local replays have a significant positive effect on place field correlations, independently of cell category (main effect of local replay *F*(1,14851) = 28.89, *p* < 0.001). Remote replays have no consistent effect on place fields (no main effect of remote replay *F*(1,15522) = 0.78, *p* = 0.39). See Table S7.

Episode representations stabilized rapidly over the first few laps (Fig.4B), on a timescale paralleling the previously reported increase in local replay rates (Fig.3A), and a decrease in decoding errors (Fig.4C). As the number of experienced episodes grew across the session, the number of newly recruited cells declined, with most cells participating in multiple episode representations (Fig.4D). Interestingly, early spiking activity on the first lap predicted subsequent recruitment into the current representation: cells with higher peak firing were more likely to become incorporated into the representation than cells with initially low firing rates (Fig.4E). The rate maps of newly recruited cells rapidly stabilized to a higher correlation value than shared cells (Fig.4F, Table S7).

We next asked whether different forms of replay preferentially recruited distinct cell categories. As expected, local replay was dominated by shared and newly recruited cells, whereas remote replay primarily recruited shared and past cells (Fig.4G). Notably, shared cells constituted the majority drivers of both replay types and participated equally in local and remote replay (Fig.4H). While shared and past cells maintained stable replay participation levels across laps, newly recruited cells progressively increased their participation in local replay, on a timescale closely matching that of map stabilization (Fig.4H). This precisely timed evolution of cell participation in local replay and map stabilization suggests that local replay may contribute directly to the stabilization of the current representation.

If local replay contributes to stabilizing the current representation by integrating newly recruited cells with those shared across representations, then replay participation – particularly for newly recruited cells - should predict an increase in rate map stability. To test this, we regressed the correlation between rate maps on consecutive laps for each cell (Ficher transformed) as a function of the number of local and remote replay events in which that cell participated (GLMM; see Methods; Fig.4I, Table S7). Rate map correlations increased with local replay participation for both shared and newly recruited cells, but did not change with remote replay participation (Fig.4J, Table S7). Contrary to part of our original hypothesis, there was no cell category × local replay interaction, with local replay equally stabilizing both cell categories. In summary, replay content is built upon a common representational scaffold, with local replay helping selectively incorporating newly recruited elements of the current episode and remote replay reactivating elements specific to past episodes in a manner that did not interfere with the current representation.

### Modeling replay as a temporally decaying strength trace

The main tenet of memory triage is for some memories to be replayed more during sleep than others. So how are sleep replay rates best predicted? So far, our analyses showed replay prioritization was unaffected by reward value during wakefulness and sleep. Sleep replay rates could not be predicted simply by the cumulative amount of prior replay as previously postulated (*32*), since the episode that accumulated the most replay events before sleep (Track 1) was not the one replayed most during subsequent sleep. Instead, replay during sleep POST was dominated by the most recent episode, mirroring the recency bias already observed during REST periods, which may reflect heightened excitability of recently active ensembles.

We conjecture that some form of temporal decay is necessary to provide a balance in replay rates, without which each new episode would need to outcompete earlier experiences, leading to a runaway escalation of replay rates. At the same time, episodes that received more replay during experience would still retain a competitive advantage. A successful model of memory triage must therefore reconcile two competing influences: the amount of replay of an episode during experience and the temporal decay of its replay probability before sleep. We therefore asked whether sleep replay rates could be predicted by a temporally weighted replay-strength trace that combines both factors.

We modeled replay strength (probability of replay) for each episode as a function of its replay history before sleep. In this model, each replay event produced an instantaneous increase in episode-specific replay strength, followed by an exponential decay over time. Thus, episodes replayed more often, and more recently, were predicted to have higher replay strength at sleep onset (Fig.5A). This model provides a simple framework for linking replay during experience and rest to subsequent replay during sleep. Importantly, the decay constant τ should *not* be interpreted as a direct biological estimate of synaptic, excitability, or neuromodulatory dynamics.

**Fig. 5:**
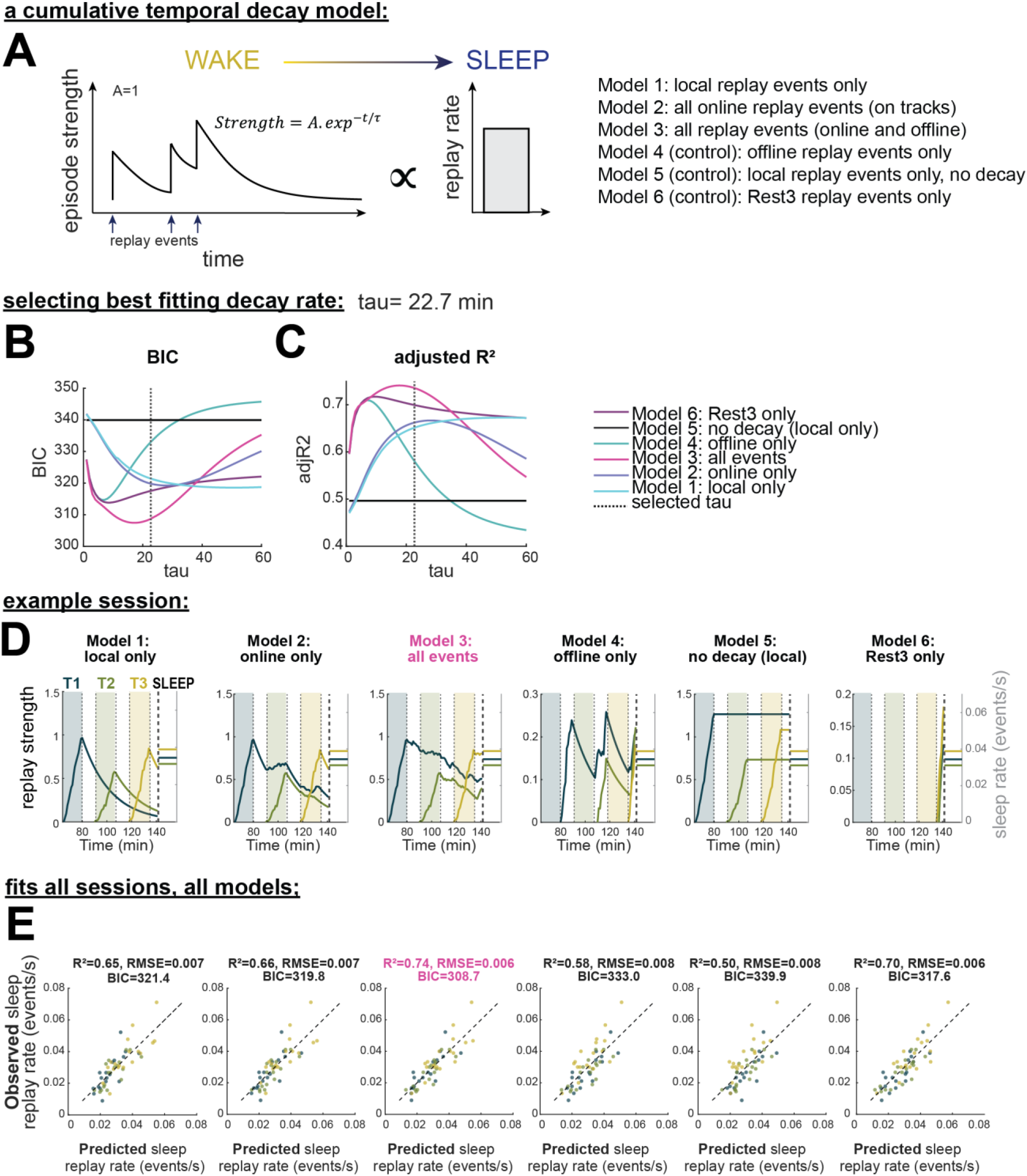
Cumulative temporal decay model of replay. **A**: schematic of the proposed model. Each replay event triggers an instantaneous increase in the replay strength - a proxy of the probability of future replay. This increase is immediately followed by an exponential decay with a fixed rate (τ). Replay events closer in time contribute to a larger replay strength than replay events spaced further in time. The replay strength prior to sleep determines initial sleep replay rates. Different sets of replay events are included to test out which contribute to building this replay strength. **B**: parameter search and selection of the decay rate τ to minimize the combined BIC across the first 3 models (excluding controls). **C**: corresponding adjusted R^2^ value for each τ value, for each model. **D**: example session (RIGEL, session 3) of the evolution of the replay strength of each episode as a function of time and replay events selected to contribute to replay strength. In colored overlay, the time periods where the animal is on a track (T1, T3 and T3). PRE sleep was not considered to contribute to replay strength and therefore is not shown. On the right vertical axis is shown the corresponding observed replay rate in the first 15min of sleep POST. A good prediction conserves the relative order and distances between sleep replay rates and the predicted replay strengths just prior. **E**: predicted vs observed sleep replay rates for each model, color coded by episode, and associated goodness of fit metrics.

We compared six variants of the model. The first three tested which replay events contribute to the strength trace: local online replay only, all online replay regardless of content, all pre-sleep replay events including online and offline periods, with the assumption that replay during the experience is a large determinant of initial episode strength. Three additional models served as controls: a local replay model without temporal decay, offline replay events only, and a model using only offline replay during the final REST period immediately preceding sleep (Fig.5A).

We selected the decay constant τ by minimizing the average Bayesian Information Criterion (BIC) across the first three models, which incorporate online replay events, testing values between 1 to 60 minutes (Fig.5B). We then visualized the resulting strength traces for individual sessions and compared the predicted relative strength of each episode at sleep onset with its observed sleep replay rate (Fig.5D). Each model is not expected to predict exact sleep rates, but rather to capture the relative allocation of sleep replay across episodes. For example, the no-decay local replay control (model 5), which corresponds to a simple “most replayed episode wins” rule, failed to capture the observed prioritization of recent episodes for that session (Fig.5D).

We regressed episode strength immediately before sleep against observed sleep replay rates and compared predicted and observed rates across models (Fig.5E). The best-performing model incorporated all replay events before sleep, followed by models using all online replay events and local online replay only. The Rest 3-only control (Model 6) also explained a large portion of the variance, as Rest 3 replay rates are highly predictive of sleep replay rates (qualitatively consistent with Fig.3E,G). It could also be partially explained by τ approximating the temporal binning of replay-rate estimates, in our case 15-minute sleep bin, even in the absence of a true temporal decay signal, which can also be observed in the BIC minima for offline only models (Models 4 and 6). However, the full all-events model (Model 3) outperformed this control across all τ values with ΔBIC_τ=22.7_ > 11, arguing against that interpretation, and suggesting that sleep replay is not determined solely by the most recent rest period. Additionally, the poor performance of the no-decay model indicates that cumulative local replay history alone is insufficient to explain sleep replay rates.

## Discussion

We investigated how competing experiences competed for access to replay before sleep and subsequent consolidation (*1–4*). We varied two key components of episodic memories, their biological utility through reward value, and their relative timing through recency. Contrary to current theories, replay prioritization was not determined by reward value but by behavioral states favorable for replay. Instead, replay was progressively redistributed across current and past experiences as additional episodes accumulated, indicating that replay is constrained by a finite budget. In this competitive setting, temporal proximity to sleep emerged as a major determinant of subsequent sleep replay rates, which can be modeled from temporally decaying replay-strength traces. Importantly, local replay stabilized emerging hippocampal representations, linking replay to both memory formation and consolidation.

### Replay-permissive behavioral states, not reward, drive replay rates

Utility-maximizing theories of replay prioritization predict that experiences with higher reward should be replayed more (*21*). These theories were recently extended by incorporating recency-weighted reward or goal history, to account for conflicting results of replay ‘lagging’ behind the current goal (*22*, *29*, *45*). We instead hypothesized that previously reported reward effects might arise because reward alters replay opportunity rather than replay probability.

Consistent with this idea, we found that replay rate and quality did not increase with reward value or reward history, despite clear behavioral evidence that rats valued and distinguished the reward conditions. This contrasts with reports of reward-related increases in replay rates (*23–26*, *46*), suggesting that such effects were driven by changes in behavior, especially increased immobility at higher-value reward sites (*23*, *26*, *42*), and therefore should be reinterpreted as alterations in replay opportunity rather than replay probability.

This reinterpretation is particularly relevant because many influential demonstrations of reward-modulated replay resulted in systematic differences in the amount of time animals spent in replay-permissive states. Singer & Frank (2009) (*46*) reported increased replay rates at rewarded compared to unrewarded goals. However, rewarded trials necessarily involved reward consumption and likely greater immobility, providing more opportunities for replay. Additionally, replay increased even before animals had fully learned the task, further indicating that reward consumption rather than reward expectation drove the effect.

Similarly, Ambrose et al. (2016) (*23*), reported that reverse but not forward replay scaled with reward value, a finding that has strongly influenced subsequent computational theories (*21*, *22*, *47*). However, reverse replay occurs preferentially later during pauses (*48*), and animals in Ambrose et al. (2016) spent longer at decreased speeds at large reward sites. Taken together with our results, these observations suggest a simple alternative: larger rewards prolonged pauses, increasing the likelihood that replay progressed into a reverse-replay state.

Generally, experimental paradigms equating or accounting for time spent at goals and reward sites across conditions are rare. Consistent with the present results, we previously demonstrated that neither place-cell activity nor a population-level goal-related signal varied with reward value when time spent at goal locations was controlled (*49*). Although replay was not directly measured, it is possible that our previous findings were an early sign that hippocampal replay-related activity is relatively insensitive to reward value itself.

This reinterpretation of behavior, rather than reward, as altering replay opportunity fundamentally challenges current theories of replay prioritization, and more broadly invites a reconsideration of replay’s proposed roles in temporal credit assignment and planning.

However, it does not fully exclude a role for reward in replay prioritization. Reward may modulate replay under conditions not captured here, such as when future task performance depends explicitly on memory retrieval, indirectly through behavioral state, or hippocampal–cortical coordination. Determining how these influences interact will be important for future studies seeking to identify the mechanisms of replay prioritization.

### Past and present episodes compete for limited replay

We also provided evidence for a rate limited replay ‘budget’ within wake bouts. As additional episodes were experienced, local replay constituted a progressively smaller fraction of replay events, whereas remote replay of previous episodes increased. This redistribution reveals a trade-off in replay between local and remote episodes, and extends previous reports that replay content is not restricted to the current environment (*6*, *50*). Evidence for a rate limited replay budget is consistent with reports that it is both costly (*15*, *16*) and homeostatically regulated (*17–19*).

What drives remote replay competition, however, remains unclear. One possibility is that remote replay emerges because episodes share a substantial fraction of their underlying neuronal representation. Cells participated in multiple episode representations, providing a natural route through which activity in the current episode could reactivate related prior experiences. Remote replay could also be prompted by the reactivation of schema-like cortical representations (*11*, *51–54*), or by prolonged increases in neuronal excitability and ensemble coordination following novel experiences (*6*, *9*, *35*, *40*). Because environment novelty was controlled in our study, it is not possible to investigate this possibility in the current dataset.

Consistent with recent theoretical work which proposes that remote replay helps prevent catastrophic forgetting (*39*), we found that remote replay did not destabilize the current representation, suggesting that replay of prior experiences is not necessarily detrimental to ongoing map formation. Whether remote replay actively protects memories from interference, however, remains an open question.

### Local replay is associated with compositional map stabilization

A finite replay budget created competition between previous and current episodes, raising the question of how replay allocation affects the representation currently being formed. Recent compositional accounts propose that hippocampal representations emerge by combining newly recruited elements with structure reused across related experiences (*38*). Consistent with this view, many cells were reused across episode representations, some of which may reflect recurring environmental features or statistical regularities (*55–57*). Newly recruited cells progressively increased their participation in local replay as representations stabilized, whereas shared cells remained active across both local and remote replay. Moreover, local, but not remote, replay predicted subsequent increases in rate map stability. These findings suggest that local replay is closely linked to the integration of newly recruited elements into a broader representational structure, and is consistent with causal evidence that optogenetic suppression of awake sharp-wave ripples impairs subsequent place-field stability (*36*).

Several additional observations connect our findings to prior work on hippocampal representation formation. Newly recruited cells stabilized more rapidly and reached higher rate map correlations than shared cells, consistent with previous reports of broadly tuned rigid and highly spatially-selective plastic components within hippocampal representations (*37*, *58*). Cells with stronger activity during initial exploration were also more likely to be incorporated into the emerging representation, consistent with activity-dependent recruitment and stabilization (*59*, *60*). Replay playing a role in map building may also explain why replay rates increase in novel environments (*32*, *35*): if replay contributes to the stabilization of newly modified representations, replay demand should scale with the amount of representational updating required.

Alternatively, replay and stabilization may both reflect representational updating driven by state prediction errors, which are expected to be greatest in the early laps of a novel experience. Under this interpretation, as the environment becomes familiar and predictable, prediction errors would decrease, representations stabilize, and replay rates diminish.

Representational updating could additionally provide a route by which reward influences replay. Rather than reflecting reward value itself, replay may track prediction errors arising from unexpected reward outcomes (*31*, *61*). If so, unpredictable changes in reward on a familiar track could increase replay despite having the same average reward value, because they require updating the animal’s internal model of the environment.

### Sleep replay is predicted by a temporally decaying episode-strength trace

Although temporal proximity to sleep is known to influence memory retention (*33*, *34*), its role in replay prioritization has received little attention. Contrary to prevailing reward-based accounts, we found that temporal proximity to sleep, but not reward, was a major determinant of elevated subsequent sleep replay. In other words, sleep replay was dominated by the most recent rather than the most replayed episode.

Replay history alone is therefore not sufficient to predict sleep replay rates. We developed a model that places temporal proximity to sleep as a fundamental feature of replay prioritization. In this model, episode-specific strength traces are strengthened by replay events but progressively decay over time. Therefore, replay does not simply accumulate. Instead, each replay event transiently increases the priority of an episode for future replay, while that priority gradually weakens unless refreshed by subsequent reactivation. Sleep replay allocation between episodes is therefore jointly predicted by how often each episode was replayed, when those replay events occurred relative to sleep, and how much competing replay intervened. Importantly, we found that all events, whether online or offline, were necessary to best predict sleep replay rates.

This finding offers an additional perspective on the function of remote replay. Although replay of previous episodes competes with replay of the current experience during wakefulness, it may simultaneously refresh the strength of older memories, allowing them to remain competitive for subsequent sleep replay. Remote replay could therefore help bridge gaps between online experience and offline consolidation.

More broadly, our proposal of replay-strength traces resonate with theories in which memory traces are maintained through repeated reactivation and decay (*3*, *62*, *63*). At the behavioral level, it provides a potential mechanistic link between replay and observations that memories encoded closer to sleep are preferentially retained (*33*, *34*).

Unlike other normative models of replay, ours does not seek to fully explain why replay occurs (*21*, *22*, *47*). Accordingly, the estimated decay time of episode-strength traces is not intended as a direct biological measure of synaptic, excitability-related, or neuromodulatory decay. Likewise, the model does not attempt to explain what generates replay rate. Influences of behavioral state, competition, cortical activity, and other factors are treated implicitly.

Instead, our model is intended to provide a working framework describing how replay history relates to subsequent sleep replay.

Overall, this study challenges the prevailing view that replay prioritization is primarily driven by utility. We demonstrate that while awake replay strengthens episodes, competition for a limited replay budget and passing time gradually erode their priority, placing temporal proximity to sleep as a key factor for consolidation. These findings call for a reassessment of existing theories of utility-based replay allocation, a revisit of previous empirical findings through the lens of replay opportunity and competition and an integration of temporal dynamics into normative accounts of memory prioritization.

## Supporting information

Statistical Summary Tables

## Acknowledgments

The authors thank Marios Panayi for discussions relating to statistical analyses; Marta-Huelin Gorriz, Lilia Kukovska, Julieta Campi and Sophie Renaudineau for technical assistance. Rat schematics in Fig.1A-B adapted with permission from SciDraw.io (https://zenodo.org/records/13371979, https://zenodo.org/records/13371977).

## Funding

Biotechnology and Biological Sciences Research Council BB/M009513/1 (MT)

Biotechnology and Biological Sciences Research Council BB/T005475/1 (DB)

European Research Council CHIME (DB)

Human Frontier Science Program RGY0067/2016 (DB)

Royal Society RG\R1\251083 (ED)

## Author contributions

Conceptualization: MT, DB. Methodology: MT, DB. Formal analysis, Investigation & Visualization: MT. Funding acquisition: MT, DB. Supervision: MT, ÉD, DB. Writing – original draft: MT. Writing - review & editing: MT, ÉD, DB

## Supplementary Materials

Materials and Methods

Supplementary Text

Figs. S1 to S6

Tables S1 to S9

