## Supplementary Materials for "Time, but not reward, shapes replay-based episodic prioritization"

**The PDF file includes:**

Materials and Methods
Supplementary Text
Figs. S1 to S6
Tables S1 to S9 (separate file)

### Materials and Methods

Animal housing and care

Five male Lister-hooded rats were implanted with microdrives. Prior to surgery, rats were kept at 90% of their free-feeding weight and housed in pairs on a 12 h light/dark cycle, with 1 h of simulated dusk/dawn. Following recovery from surgery, animals were single-housed and food restricted, with their weight and health checked daily. All experimental procedures and postoperative care were approved and carried out in accordance with the UK Home Office, subject to the restrictions and provisions contained within the Animal (Scientific Procedures) Act of 1986.

### Methods

Methods are largely identical to those described in (Tirole et al., 2022). All analyses were performed using custom scripts in MATLAB and R.

Surgery

Animals were deeply anesthetized with isoflurane (1.5–3% at 2 L/min) and implanted with custom-made microdrive arrays carrying a total of 16-24 independently moveable tetrodes (modified from microdrive published by (Davidson et al., 2009), and 'poor lady drives' from Axona Ltd). Each tetrode consisted of a twisted bundle of four tungsten microwires (12 or 17 µm diameter, Tungsten 99.95% CS, California Fine Wire), gold-plated to reduce impedance to <200 kΩ. The microdrives targeted dorsal hippocampal CA1 bilaterally (AP: –3.48 mm; ML: ±2.4 mm from Bregma). After surgery, animals were housed individually and allowed to recover with food and water ad libitum for a week before returning to being kept at 90% of their pre-operative free-feeding weight.

Reward preference test

Following recovery from surgery, but prior to start of recordings, we ran a reward preference assay. Animals were placed on a (20 x 80) cm platform with two ceramic bowls filled with 15mL of undiluted or 1:1 dilution of chocolate flavored soy milk. Each trial consisted of 2 minutes where the animal could freely sample the liquids, after which the animal was removed from the platform, and the remaining amount of liquid measured. The bowls were then refilled and placed back in a pseudo-random manner at either end of the platform. 12 trials were performed by each animal. The rats were therefore familiar with both rewards prior to the start of recordings.

Experimental design
Each session began with a 1 h rest period, during which rats were placed in a black circular enclosure 20 cm in diameter, surrounded by a 50 cm-tall view-obstructing plastic sheet, hereafter referred to as the rest pot. Rats were then exposed sequentially to three novel linear tracks with identical geometry and dimensions. On each track, rats were allowed to run back and forth for 15 min and received liquid rewards, 0.1 mL (delivered by infusion pumps, dual Aladdin, WPI), at both ends when they correctly alternated between reward sites. Custom Bonsai (Lopes, G., & Monteiro, P. 2021) and Arduino scripts controlled the task logic and reward delivery system. Between track exposures, rats were returned to the rest pot for approximately 10 min (REST 1 and REST 2). After exposure to the third track, rats were returned to the rest pot for a final rest period lasting approximately 1.5 to 2 h.

To create novel and distinct track episodes, the tracks' geometry, global cues, and local cues were changed daily. Global cues consisted of wall-mounted panels, whereas local cues consisted of textured fabrics placed on the tracks. Wall dividers were positioned between tracks to create sub-rooms within the recording room. When necessary to maintain a high single-unit yield, tetrodes were adjusted after the end of the recording session.

| Day | Track 1 | Track 2 | Track 3 |
|---|---|---|---|
| 1 | LOW | LOW | HIGH |
| 2 | LOW | HIGH | LOW |
| 3 | HIGH | LOW | LOW |
| 4 | HIGH | HIGH | LOW |
| 5 | HIGH | LOW | HIGH |
| 6 | LOW | HIGH | HIGH |

Table showing all conditions recorded. For this study, all animals experienced Days 1-3 block first, followed by Days 4-6, but with pseudo randomized session within these blocks:

| Day | Track 1 | Track 2 | Track 3 |
|---|---|---|---|
| 1 | LOW | LOW | HIGH |
| 2 | HIGH | LOW | LOW |
| 3 | LOW | HIGH | LOW |
| 4 | HIGH | LOW | HIGH |
| 5 | HIGH | HIGH | LOW |
| 6 | LOW | HIGH | HIGH |

Example: session counterbalancing for Polaris.

Before recordings began, rats were trained for approximately 2 days, 30 min per day, in a separate room to run back and forth on a linear track to collect liquid rewards at each end. For rat Navi, two condition sequences, low/low/high and high/high/low, were each repeated once due to human error.

Recordings
Data were collected using a digital Neuralynx acquisition system (Digital Lynx SX, Neuralynx). Neural signals, referenced to a screw implanted above the animal's cerebellum, were digitized at the headstage. The continuous raw electrophysiological signal was sampled at 30 kHz, highpass filtered > 0.1 Hz, and stored for subsequent processing. The animal's behavior was recorded by

an overhead camera. The 25Hz video feed was synchronized by the system to the neural signal, and the animal's position was tracked using green and red LEDs mounted on the headstage.

Spike detection and unit isolation

Spike data were extracted using the semi-automated clustering software KlustaKwik 2.0 (K. Harris, http://klustakwik.sourceforge.net/) and manually curated in Klustaviewa (https://github.com/klusta-team/klustaviewa). Putative single units were identified based on spike waveform characteristics, clean inter-spike interval distributions, and stability across the recording session. Remaining clustered activity was classified as either multi-unit activity (MUA) or noise.

LFP processing

The power spectral density of the Local Field Potential (LFP) was estimated using Welch's method (MATLAB's *pwelch*) to identify channels with the strongest delta, 1-4 Hz; theta, 4-12 Hz; and ripple, 125-300 Hz, power, as well as the channel showing the largest difference between normalized theta and ripple power. LFP signals from the selected channels were then downsampled from 30 kHz to 1 kHz and band-pass filtered (MATLAB's *filtfilt*). Instantaneous phases were estimated using the Hilbert transform.

Place cell selection and rate map calculation

Putative principal cells were defined as single units with a half-width at half-maximum > 500 µs and a mean firing rate < 5 Hz across the full recording session. For place-cell classification, position data were discretized into 10 cm bins (20 position bins for a 2m track), spike trains were filtered to include only spikes emitted when animals were running between 5 and 50 $cm.s^{-1}$. A principal cell was classified as a place cell if its rate map had a peak firing rate > 1 Hz. Place cells were additionally required to show stable spatial firing, defined as a peak firing rate > 1 Hz in both the first and second halves of each track exposure.

For directional fields, spiking and position data were separated by running direction on each track, and the place cell selection and rate map calculation procedures described above were repeated for each direction and track.

Candidate replay events detection

MUA was binned in 1 ms steps and smoothed with a Gaussian kernel, $\sigma = 5$ ms. Bursts were initially selected if their z-scored activity exceeded 3 and their maximum duration was 300 ms. Events passing this threshold were then filtered according to behavioral and spiking criteria: candidate events were retained only when the animal's speed was below 5 cm $s^{-1}$, at least five different units were active, and event duration was between 100 and 750 ms, corresponding to at least five consecutive 20 ms bins. Events occurring within 50 ms of one another were merged. Ripple power was also used as an additional criterion: the ripple-band-filtered LFP was smoothed with a 0.1 s moving-average filter, z-scored, and required to exceed a threshold of 3. To improve replay detection and avoid excluding events with noisy probability decoding at the beginning or end, candidate replay events were also additionally analyzed after splitting them into two sub-events. The split point was defined as the time of minimum MUA activity within the middle third of the event, providing a natural midpoint. Both resulting segments were decoded and tested for significance independently using the same criteria as intact candidate events, including the minimum-duration requirement, but with an adjusted significance threshold of $p < 0.025$.

For candidate replay event rates presented in Figures 2 and 3, the criterion of at least 5 single units active was removed to limit biasing results toward track replays.

Bayesian decoding

A naïve Bayesian decoder was used to reconstruct the animal's spatial trajectory during both behavior and candidate replay events from CA1 hippocampal spiking activity (Zhang et al.,1998). Since a naïve Bayesian decoder was used, the prior was assumed to be uniform across positions and set to 1. The normalization constant was defined as the summed posterior probability across all tracks. Position was decoded using 250 ms time bins during behavior and 20 ms time bins during replay events. All place cells with a place field on at least one track were included in the decoder.

Decoding error was calculated as the absolute difference between the animal's true position and the maximum-likelihood decoded position.

Replay scoring

Replay events were scored using the weighted correlation between decoded posterior probabilities across position and time. The statistical significance of each candidate replay event was assessed by comparing its weighted correlation score with three shuffle distributions, each generated from 1,000 iterations.

1. spike train circular shift shuffle: spike-count vectors from each cell were independently circularly shifted in time within each replay event before decoding.
2. place field shift shuffle, each rate map was circularly shifted in space by a random number of position bins before decoding.
3. posterior circular shift shuffle, posterior probability vectors from each time bin were independently circularly shifted in the position axis by a random amount.

Candidate events were considered significant only if their weighted correlation score exceeded the 95th percentile of all three shuffle distributions. In a small number of cases, replay events were significant for both tracks. These events were assigned to a single track using a Bayesian bias score, calculated separately for each track as the sum of the posterior probability matrix for that track normalized by the total posterior probability summed across both tracks. Events were assigned to a given track only when the corresponding Bayesian bias score exceeded 60%; otherwise, they were excluded from further analysis.

To classify forward and reverse replay, the same procedures were repeated using directional rate maps. The slope of each replay event was estimated using a line-fitting procedure (Ólafsdóttir et al., 2016). A replay event was classified as forward when decoded content corresponded to direction 1 with a positive slope or direction 2 with a negative slope. A replay event was classified as reverse when decoded content corresponded to direction 1 with a negative slope or direction 2 with a positive slope.

State detection during PRE and POST

State detection was performed using position and LFP data restricted to periods when the animal was in the rest pot. Speed was smoothed with a 30 s moving-average window and z-scored. Immobility was defined as periods in which z-scored speed remained below −0.5 for at least 10 s. Delta and theta power were smoothed with a 15 s moving-average window. Theta/delta measures were then computed as both the theta/delta envelope ratio and the normalized theta–delta preference, defined as $(theta - delta)/(theta + delta)$. Each time point was classified as wake, quiet rest, NREM, or REM:

1. NREM was defined as immobile periods with a normalized theta-delta preference ≤ −0.1 and a theta/delta z-score ≤ 0.8, with a minimum bout duration of 5 s.
2. REM was defined as immobile periods with a normalized theta–delta preference ≥ 0.1 and a theta/delta z-score > 0.8, with a minimum bout duration of 2 s.
3. Quiet rest was defined as periods with z-scored speed ≤ 0 that were not classified as NREM or REM.
4. Wake was defined as periods of mobility z-scored speed > 0.

NREM bouts shorter than 5 s and REM bouts shorter than 2 s were reassigned to the surrounding state when flanked by identical states on both sides (e.g. too brief NREM bout surrounded by Wake); otherwise, they were classified as quiet rest.

Population vector analysis (map stabilization)

To calculate the stabilization of hippocampal maps with experience of each track, lap-specific rate maps were calculated. Then, for each session, the rate maps were stacked into a 20 position bins × N cells matrix for each track and lap. The linearized population vector of rates at each position bin was then correlated with its counterpart vector obtained from the subsequent lap. The mean Pearson's correlation value was first averaged across position bins and then across conditions.

GLMMS

Unless otherwise stated, all generalized linear mixed models were run in R (afex::mixed). Data tables were imported into R, and *rat*, *track* (episode), *reward*, and *session* were coded as categorical factors. Sum-to-zero contrasts were applied to *reward*, *track*, and *session*. For each dependent variable, models included fixed effects (e.g. reward, track, and their interaction), with nested random intercepts for session within rat. For each fitted model, we extracted ANOVA results for the fixed effects, estimated marginal means (e.g.: for reward, track, and reward within track), as well as pairwise contrasts (e.g. for reward, track, simple reward effects within each track, and reward-by-track interaction contrasts). Sidak-adjusted p-values and confidence intervals were calculated using an adjusted alpha level, defined as (Maxwell & Delaney, 1990):

$$1 - (1 - 0.05 \times number\ of\ contrast\ families)^{1/number\ of\ contrasts}$$

Estimated marginal means, contrast statistics, and ANOVA tables were saved for each model, and relevant statistics are summarized in Tables S1–S9.

List of GLMMs

| formula | Dependent variable |
| --- | --- |
| Behavior measures – Table S1 | |
| ~ reward × track + (1 \| rat/session) | Total number of laps per episode |
| ~ reward × track + (1 \| rat/session) | Total time spent immobile (< 5 $cm.s^{-1}$) anywhere on the track (s), per lap |
| ~ reward × track + (1 \| rat/session) | Total time spent immobile (< 5 $cm.s^{-1}$) inside the reward zones (s), per lap |
| ~ reward × track + (1 \| rat/session) | Total time spent immobile (< 5 $cm.s^{-1}$) outside the reward zones (s), over entire episode, not per lap because too infrequent |
| ~ reward × track + (1 \| rat/session) | Median running speed of animal (> 5 $cm.s^{-1}$), per lap |
| ~ reward × track + (1 \| rat/session) | Median sub-threshold speed of animal (< 5 $cm.s^{-1}$), per lap |
| Behavior measures – Table S2 | |
| ~ reward × track + (1 \| rat/session) | Total time spent immobile (< 2 $cm.s^{-1}$) anywhere on the track (s), per lap |
| ~ reward × track + (1 \| rat/session) | Total time spent immobile (< 2 $cm.s^{-1}$) inside the reward zones (s), per lap |
| ~ reward × track + (1 \| rat/session) | Total time spent immobile (< 2 $cm.s^{-1}$) outside the reward zones (s), over entire episode, not per lap because too infrequent |
| ~ reward × track + (1 \| rat/session) | Median running speed of animal (> 2 $cm.s^{-1}$), per lap |
| ~ reward × track + (1 \| rat/session) | Median sub-threshold speed of animal (< 2 $cm.s^{-1}$), per lap |
| Behavior measures – Table S3 | |
| Same as Table S1 for only three rats included in replay analyses | |
| Replay measures – Table S4 | |
| Event Rate ~ speed + lap number + (1 \| rat/session) | candidate replay events on tracks, per lap |
| Event Rate ~ speed + lap number + (1 \| rat/session) | local replay, per lap |
| Event Rate ~ speed + lap number + (1 \| rat/session) | local forward replay, per lap |
| Event Rate ~ speed + lap number + (1 \| rat/session) | local reverse replay, per lap |
| Event Rate ~ reward × track + (1 \| rat/session) | replay during PRE |
| Event Rate ~ reward × track + (1 \| rat/session) | replay during sleep PRE |

| | |
|---|---|
| Event Rate ~ reward × track + speed + lap number + (1 \| rat/session) | candidate replay events on tracks, per lap |
| Event Rate ~ reward × track + speed + lap number + (1 \| rat/session) | local replay, per lap |
| Event Rate ~ reward × track + speed + lap number + (1 \| rat/session) | local forward replay, per lap |
| Event Rate ~ reward × track + speed + lap number + (1 \| rat/session) | local reverse replay, per lap |
| Event Rate ~ reward × track + (1 \| rat/session) | replay during REST1 |
| Event Rate ~ reward × track + (1 \| rat/session) | replay during REST2 |
| Event Rate ~ reward × track + (1 \| rat/session) | replay during REST3 |
| Event Rate ~ reward × track + (1 \| rat/session) | replay during POST |
| Event Rate ~ reward × track + (1 \| rat/session) | replay during sleep POST (first 15 minutes) |
| Event ~ reward × track + (1 \| rat/session) | replay during sleep POST (overall) |
| Replay measures – Table S5 | |
| Event Rate ~ reward × track + speed + lap number + (1 \| rat/session) | local replay (at reward sites) |
| Event Rate ~ reward × track + speed + lap number + (1 \| rat/session) | local forward replay (at reward sites) |
| Event Rate ~ reward × track + speed + lap number + (1 \| rat/session) | local reverse replay (at reward sites) |
| Event Rate ~ reward × track + speed + lap number + (1 \| rat/session) | candidate replay events (at reward sites) |
| Replay measures – Table S6 | |
| ~ reward × track + (1 \| rat/session) | local replay quality (weighted correlation) |
| ~ reward × track + (1 \| rat/session) | local forward replay quality (weighted correlation) |
| ~ reward × track + (1 \| rat/session) | local reverse replay quality (weighted correlation) |
| Building maps - Table S7 | |
| fieldCorr_z ~ track + cell category×lap number (centered) + cell category×number of local replays cell was active in (centered) + number of remote replays cell was active in (centered) + (track + lap number (centered) + cell category \|\| cell id) | Fisher transformed ratemap correlation of each cell between consecutive laps |
| Replay, extension quiescence and sleep - Table S8 | |
| ~ sub threshold speed (*fitlm*) | local replay rate |
| ~ 1 + zscored speed + (1 \| rat) + (1 \| rat:session) (*fitlme*) | Replay rate during quiet states in POST |
| ~ 1 + cumulative time in sleep + (1 \| rat) + (1 \| rat:session) (*fitlme*) | Replay rates over sleep POST |

| Replay and reward history - Table S9 | |
|---|---|
| ~ reward + speed + lap number + (1 \| session) | Local replay rates on T1 (random effect of rat led to singularities as it didn't explain any variance) |
| ~ reward × previous reward + speed + lap number + (1 \| rat/session) | Local replay rates on T2 |
| ~ reward × previous reward × track + speed + lap number + (1 \| rat/session) | Local replay rates on T2 & T3 |

Note, in MATLAB, (1 | rat) + (1 | rat:session) is equivalent to (1|rat/session) in R

<u>Cell contribution replay</u>

Cell category participation across replay types: for each half-lap segment containing one reward-site visit, cells were classified as active in a replay event if they emitted at least one spike during the replay window. For each cell, we calculated the proportion of local or remote replay events in which it was active, then averaged these values within predefined cell categories.

Percent involvement of cell categories in replay events: for each replay event within each half-lap segment, we calculated the proportion of active cells belonging to each predefined cell category, defined as the number of active cells in that category divided by the total number of active cells in the event. These event-level proportions were averaged across replay events within each replay type and half-lap segment.

<u>Temporal decay model</u>

We estimated episode-specific replay strength as a function of each episode's replay history prior to sleep. In the model, every replay event caused an immediate increase in replay strength for that episode, which then decayed exponentially over time. Replay strength at each time point was calculated by applying an exponential decay kernel to the binned replay-event counts:

$$S_i(t,\tau) = \sum_t C_{i,t}\, exp(-\frac{T_{sleep} - t}{\tau})$$

where $C_{i,t}$ is the binned count of replay events for episode i at time bin $t$, $T_{\text{sleep}}$is the onset time of sleep POST, and τ is the decay time constant.

Several alternative replay-strength models were evaluated by varying the event-count term, $C_{i,t}$. These included models based on local online replay only, all online replay events irrespective of content, all pre-sleep replay events across both online and offline epochs, and offline replay events alone. Two control models were also included: one based on local replay counts without temporal decay, and another using only offline replay from the final REST epoch immediately before sleep (Rest 3).

For each candidate τ, replay strength was used to predict the observed replay rate using a linear mixed-effects model (*fitlme*):

$$\text{SleepReplayRate} \sim \text{Strength at sleep onset} + (\text{session} \mid \text{rat}).$$

Model performance was evaluated across a range of τ values (1min to 60min) using adjusted $R^2$, Root Mean Squared Error (RMSE) and Bayesian Information Criterion (BIC). A single global τ was selected by minimizing the summed BIC across the main candidate models.

$$RMSE: \sqrt{(observed\ rate\ -\ predicted\ rate)^2}$$

The same analysis pipeline was also run with models trained against REST3 replay rates rather than sleep replay rates; these REST3-trained models were then evaluated/tested against post-task SLEEP replay rates.

### Supplementary Text

Extended description of behavior and statistical interpretation (Table S1)

Reward significantly influenced several behavioral measures:

Rats completed an average of 14 ± 5 laps per track, corresponding to an average of approximately 28 rewards per track. Rats ran significantly more laps for HIGH than LOW rewarded tracks (estimated mean LOW= 12.11 laps, HIGH= 15.85 laps. For the number of laps completed within an episode, there was a significant main effect of Reward ($F_{1,60.220}=22.479$, $p < 0.001$) and a significant Reward × Track interaction ($F_{2,78.380}=3.358$, $p=0.040$, but no main effect of Track ($F_{2,54.731} = 2.014,\ p = 0.143$. Pairwise contrasts revealed that this interaction was primarily driven by T3, where animals completed significantly more laps in the high-reward condition (T3(HIGH-LOW) $\beta = 6.005,\ t = 3.967,\ p < 0.001$), whereas reward effects were not significant within T1 or T2 after correction ($\alpha = 0.016$).

Despite this increased running, the rats did not consistently spend more time at the reward sites for HIGH vs LOW. Rats spent an average of 12s at each reward site (estimated means LOW=13.41s, HIGH= 14.62s, for significant simple effect T2: LOW= 12.93s, HIGH= 17.34s). Time spent immobile at reward sites showed a significant Reward × Track interaction ($F_{2,643.072}=4.252$, $p=0.015$), with no main effects of Reward ($F_{1,1203.696} = 3.048,\ p = 0.081$) and a trend for Track ($F_{2,1210.291} = 2.979,\ p = 0.051$). However, no individual pairwise contrasts survived correction ($\alpha = 0.016$), suggesting that the interaction reflected relatively subtle distributed effects rather than a strong effect localized to a single track.

The rats' subthreshold speed ($v < 5$ cm.s$^{-1}$) did vary with reward value corresponding to different levels of stillness: most likely between grooming, consuming reward, looking around, etc. Rats were more immobile when consuming the HIGH reward (estimated mean LOW= 2.07 cm.s$^{-1}$, HIGH= 1.72 cm.s$^{-1}$). Subthreshold speed was strongly modulated by Reward ($F_{1,1218.671}=89.628$, $p < 0.001$), with no main effect of Ttrack ($F_{2,1205.749} = 0.047,\ p = 0.955$) and no Reward × Track interaction ($F_{2,819.191} = 0.909,\ p = 0.403$). Reward effects were most pronounced in T3(HIGH-LOW) ($\beta = -0.348,\ t = -4.868,\ p < 0.001$), whereas effects in T1 and T2 were not significant.

Running speed (v > 5 cm.s$^{-1}$) was similarly modulated by Reward ($F_{1,1216.228} = 45.064,\ p < 0.001$) and showed a significant Reward × Track interaction ($F_{2,1209.363} = 12.704,\ p < 0.001$), with only a trend toward a main effect of Track ($F_{2,1202.306} = 2.782,\ p = 0.062$). Reward-dependent changes in running speed were strongest in T3 ($\beta = 2.858,\ t = 7.181,\ p < 0.001$), whereas effects in T1 and T2 did not survive correction.

We also observed differences in immobility away from reward sites. Rats spent more time immobile (anywhere on the track) for LOW than HIGH. Total time spent immobile anywhere showed a significant main effect of Reward ($F_{1,1223.295} = 18.118,\ p < 0.001$) and a strong Reward × Track interaction ($F_{2,883.219} = 13.514,\ p < 0.001$), but no main effect of Track ($F_{2,1206.894}=0.346$, p=0.707). This interaction was driven primarily by T3, where reward strongly altered immobility T3(HIGH-LOW) ($\beta = -18.386,\ t = -6.154,\ p < 0.001$), with lower sub-threshold speed, ran faster and completed more laps (see above).

Time spent immobile away from reward sites also showed a significant effect of Reward ($F_{1,67.116} = 13.468,\ p < 0.001$), but no main effect of Track ($F_{2,55.666} = 0.339,\ p = 0.714$) and no Reward × Track interaction ($F_{2,75.439} = 1.208,\ p = 0.304$). Again, the clearest reward-related effect was observed in T3(HIGH-LOW) ($\beta = -6.583,\ t = -3.279,\ p = 0.002$).

<u>Extended description of behavior (3 replay rats) and statistical interpretation (Table S3)</u>

Reward significantly influenced behavior, leading to similar results as above:

The number of laps completed showed a significant main effect of Reward ($F_{1,36.114} = 8.502,\ p = 0.006$), but no main effect of Track ($F_{2,31.204} = 0.479,\ p = 0.624$) and no Reward × Track interaction ($F_{2,45.749} = 1.886,\ p = 0.163$). Pairwise contrasts between tracks were all non-significant. Reward effects were strongest in T3(HIGH-LOW) ($\beta = 5.058$, $t = 2.622$, $p = 0.012$), whereas effects in T1 and T2 did not survive correction.

Time spent immobile at reward sites did not show significant main effects of Reward ($F_{1,695.508} = 4.615,\ p = 0.032$), Track ($F_{2,692.986} = 2.402,\ p = 0.091$), and there was no Reward × Track interaction ($F_{2,328.171} = 2.095,\ p = 0.125$).

Subthreshold speed (v < 5 cm.s$^{-1}$) was strongly modulated by Reward ($F_{1,700.999} = 38.046,\ p < 0.001$), but not by Track ($F_{2,690.298} = 0.415,\ p = 0.661$) and showed no Reward × Track interaction ($F_{2,488.157} = 1.292,\ p = 0.276$). Reward significantly reduced subthreshold speed overall (HIGH-LOW $\beta = -0.306,\ t = -6.149,\ p < 0.001$), with significant effects in both T2(HIGH-LOW) $\beta = -0.436,\ t = -4.414,\ p < 0.001$) and T3(HIGH-LOW) $\beta = -0.250,\ t = -2.539,\ p = 0.011$), but not T1 (trend, did not survive correction).

Running speed (v > 5 cm.s$^{-1}$) showed a robust main effect of Reward ($F_{1,694.973} = 13.151,\ p < 0.001$) and a significant Reward × Track interaction ($F_{2,694.703} = 15.438,\ p < 0.001$), but no effect of Track alone ($F_{2,688.310} = 0.478,\ p = 0.620$). Reward significantly increased running speed overall (HIGH-LOW $\beta = 0.995,\ t = 3.623,\ p < 0.001$), with the strongest effect in T3 ($\beta = 3.429,\ t = 6.137,\ p < 0.001$).

Total time spent immobile anywhere showed a significant main effect of reward ($F_{1,702.311} = 7.120,\ p = 0.008$) and a strong Reward × Track interaction ($F_{2,469.019} = 9.501,\ p < 0.001$), but no effect of track alone ($F_{2,691.111} = 0.372,\ p = 0.689$). Reward significantly reduced overall immobility ($\beta = -5.770,\ t = -2.659,\ p = 0.008$), an effect driven almost entirely by T3 ($\beta = -21.871,\ t = -5.103,\ p < 0.001$), with no significant effects in T1 or T2.

Time spent immobile away from reward sites similarly showed a significant main effect of reward ($F_{1,45.988} = 6.886,\ p = 0.012$), but no effect of track ($F_{2,45.990} = 0.580,\ p = 0.564$) and no Reward × Track interaction ($F_{2,46.523} = 1.347,\ p = 0.270$). Reward significantly reduced immobility away from reward sites overall ($\beta = -3.684,\ t = -2.570,\ p = 0.014$), again primarily on T3(HIGH-LOW) ($\beta = -6.654,\ t = -2.647,\ p = 0.011$).

Overall, the behavioral patterns were highly similar to those observed in dataset comprising all animals. Reward increased the number of laps and running speed, reduced subthreshold speed, while leaving time spent immobile constant. Track identity alone contributed relatively little variance. Analyses of both datasets converge on the same central conclusion: reward strongly alters behavioral state.

<u>Extended description of replay rates x reward value and statistical interpretation (Table S4, candidate events and local replay rates)</u>

For candidate replay events detected on the tracks, there was no main effect of Reward ($F_{1,669.704} = 0.031,\ p = 0.859$), but there was a strong main effect of Track ($F_{2,693.616} = 10.304,\ p < 0.001$). The Reward × Track interaction was not significant ($F_{2,143.003} = 1.475,\ p = 0.232$). Importantly, both subthreshold speed ($F_{1,637.748} = 53.415,\ p < 0.001$) and lap number ($F_{1,680.347} = 74.441,\ p < 0.001$) were strong predictors of candidate replay event occurrence. Pairwise contrasts showed significantly fewer candidate events in T1 relative to both T2 ($\beta = -0.033,\ t = -3.291,\ p = 0.001$) and T3 ($\beta = -0.044,\ t = -4.365,\ p < 0.001$), whereas T2 and T3 did not differ ($p = 0.308$). There was no significant effect of reward overall ($\beta = 0.002,\ p = 0.860$) or within individual tracks after correction.

For local replay events, there was similarly no main effect of Reward ($F_{1,700.747} = 0.151,\ p = 0.698$), and a significant main effect of Track ($F_{2,691.591} = 3.792,\ p = 0.023$). The Reward × Track interaction was significant ($F_{2,354.616} = 3.110,\ p = 0.046$). Both subthreshold speed ($F_{1,583.952} = 26.043,\ p < 0.001$) and lap number ($F_{1,657.559} = 33.819,\ p < 0.001$) significantly predicted local replay occurrence. Pairwise contrasts revealed significantly greater local replay in T1 compared to T3 ($\beta = 0.014,\ t = 2.454,\ p = 0.014$), a trend compared to T2 ($\beta = 0.013,\ t = 3.225,\ p = 0.02,\ alpha = 0.016$) whereas T2 and T3 did not differ ($p = 0.928$). Reward did not significantly alter local replay overall ($\beta = -0.002,\ p = 0.699$) or within any individual track after correction. The Reward × Track interaction is due to a higher-order interaction effect we do not consider to reflect a systematic increase in rates by reward.

Extended description of replay rates x reward history and statistical interpretation (Table S9, local replay rates)

To investigate how replay allocation might be influenced by reward history rather than absolute reward value, where replay rates reflect a relative comparison between current and previously experienced rewards we started by replicating our previous analyses, limiting the data to Track1, where no prior reward history was available. There was no effect of Reward $F_{1,217.89} = 1.27, p = 0.26$. We then examined local replay exclusively on Track 2, where animals could potentially evaluate the current reward relative to the preceding episode. There was no effect of Reward ($F_{1,12.95} = 0.0008, p = 0.97$), of Previous Reward ( $F_{1\ ,12.38} = 1.1783, p = 0.298$), nor their interaction ($F(1{,}14.46) = 0.259,\ p = 0.618$ ). Finally, we obtained similar results when extending the analysis across Tracks 2 and 3: no effect of Reward ( $F_{1,46.618} = 0.5709\ =, p = 0.453$ ), of Previous Reward ( $F_{1\ ,54.49} = 3.331, p = 0.073$), nor their interaction ( $F_{1,399.308} = 0.2177, p = 0.641$), no other interactions, including with track were significant. Across all of these analyses, consistent with prior results, sub threshold speed and lap number significantly modulated replay rates (see Table S9).

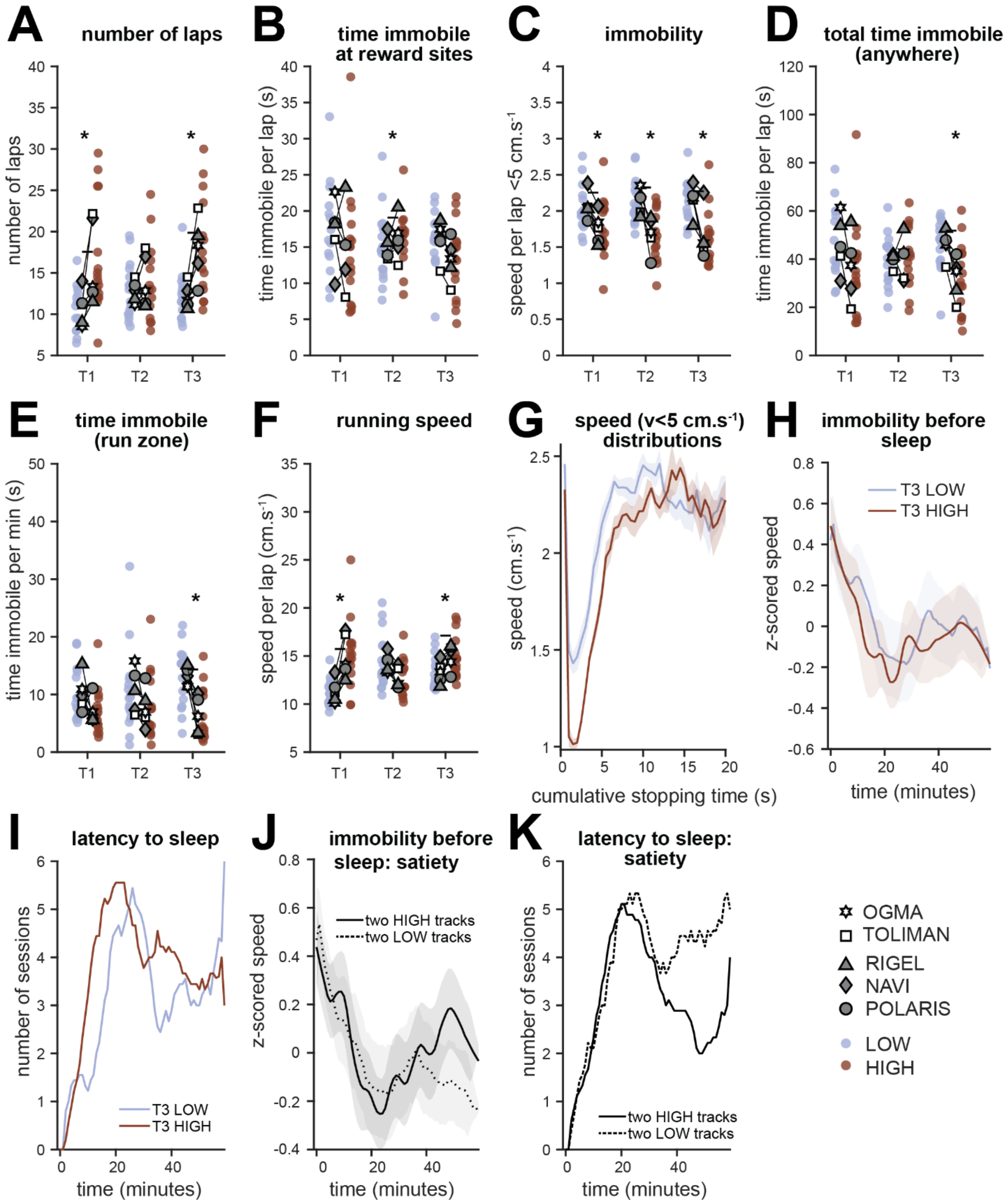

**Fig. S1. behavioral measures.**

**A-F**: Inside markers: rat averages for each measure, split for reward × track identity. Outside round markers are individual datapoints. In brown, HIGH value tracks, in blue LOW value tracks. Rats used for replay analyses have filled in grey symbols. Denoted by (*) significance of simple effects from GLMM ~ reward × track + (1 | rat/session), see corresponding **Table S1**

**and supplementary text**. **A**: number of laps on each track. **(**main effect of reward $F(1{,}60.22) = 22.48, p < 0.001$**;** reward x track interaction $F(2{,}78.38) = 3.36,\ p\ = 0.04$**;** T1(HIGH>LOW) p = 0.003; T3 (HIGH>LOW) p = < 0.001) **B**: time spent immobile at each reward site, in seconds, per lap (reward x track interaction $F(2{,}643.07) = 4.25$, p= 0.015; T2(HIGH>LOW) p = 0.001). **C**: mean sub-threshold speed (v < 5 $cm.s^{-1}$) per lap (main effect of reward $F(1{,}1218.67) = 89.63, p\ <\ 0.001$; all within reward contrasts HIGH<LOW p < 0.001). **D**: time spent immobile anywhere on the track, in seconds, per lap (main effect of reward $F(1{,}1233.29) =\ 18.11, p\ <\ 0.001$; reward x track interaction $F(2{,}883.21) =\ 13.51$, p < 0.001); T3(HIGH<LOW) p < 0.001). **E**: time spent immobile away from reward sites, in seconds, per lap (main effect of reward $F(1{,}67.11) =\ 13.46,\ p < 0.001$; T3(HIGH<LOW) p = 0.002). **F**: mean running speed (v > 5 $cm.s^{-1}$) per lap (main effect of reward $F(1{,}1216.22) = 45.06, p\ <\ 0.001$; reward x track interaction $F(2{,}1209.36) = 12.7, p < 0.001$; T1(HIGH>LOW) p = 0.001, T3(HIGH>LOW) p < 0.001). **G**: mean + sem speed distributions, in $cm.s^{-1}$, over cumulative stopping time, in seconds, split for LOW and HIGH reward sites. **H-K** data from three rats used for replay analyses. **H-I:** HIGH reward on track 3 is associated to a shorter latency to sleep and increased immobility **H**: speed distributions before sleep during POST, split depending on whether Track3 was LOW or HIGH. **I**: number of sessions where the rat has fallen asleep, as a function of time, in seconds, since the start of POST. Split depending on whether Track3 was LOW or HIGH. **J-K**: same as H and I respectively, this time split based on the number of HIGH rewarded tracks in a session (one or two) which loosely correlates with satiety, there is no difference between the curves in the first few minutes, suggesting that differences seen in H,I are not linked to overall reward value consumed.

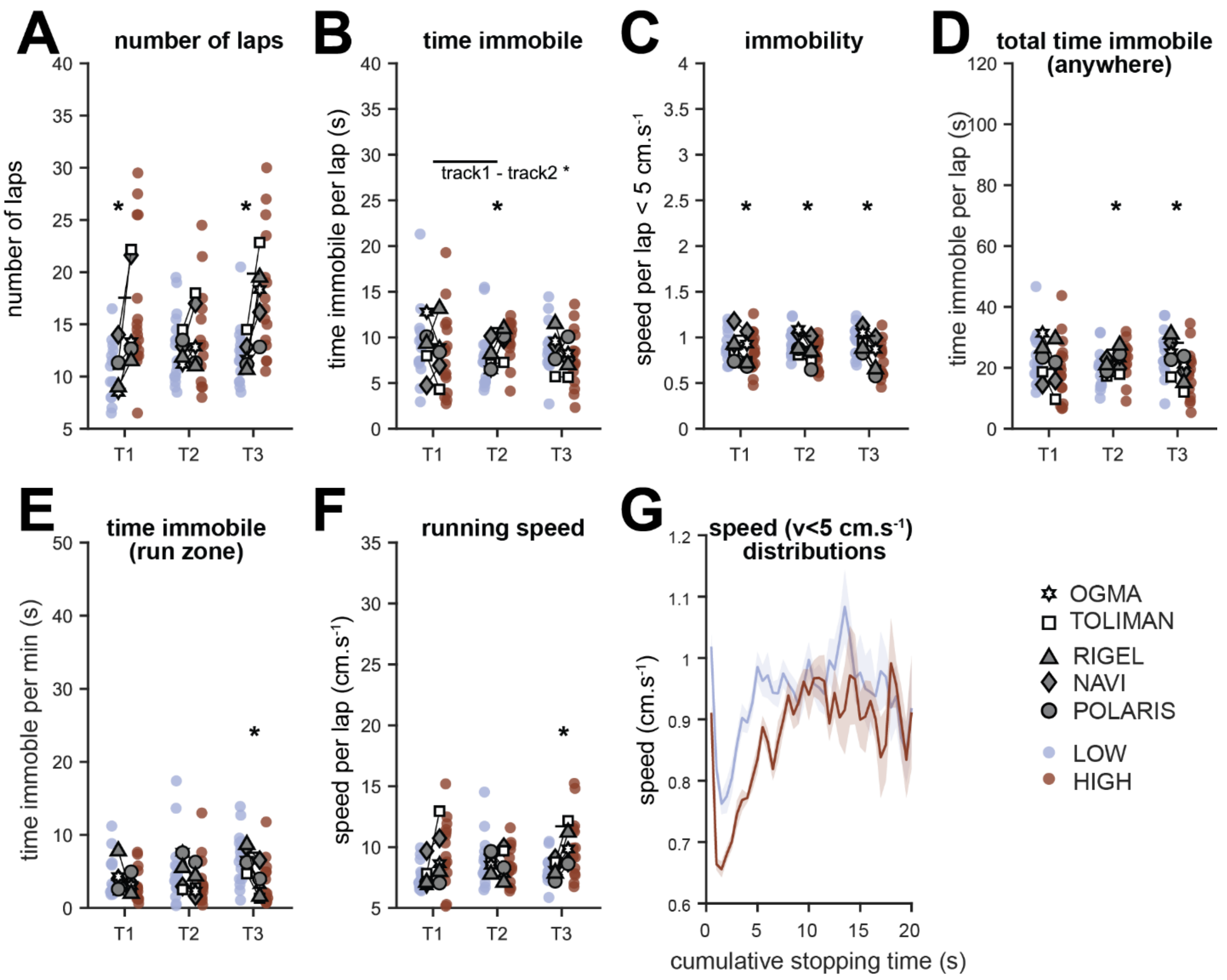

**Fig. S2. Behavioral measures when immobility speed threshold is lowered to 2cm.s$^{-1}$**

**A-F**: Inside markers: rat averages for each measure, split for reward × track identity. Outside round markers are individual datapoints. In brown, HIGH value tracks, in blue LOW value tracks. Rats used for replay analyses have filled in grey symbols. Denoted by (*) significance of simple effects from GLMM ~ reward × track + (1 | rat/session), see corresponding **Table S2**. **A**: number of laps on each track (main effect of reward $F(1{,}60.22) = 22.47, p < 0.001$); reward x track interaction $F(2{,}78.38) = 3.35, p = 0.04$); T1(HIGH>LOW) p =0.003, T3(HIGH>LOW) p<0.001). **B**: time spent immobile at each reward site, in seconds, per lap (main effect of reward $F(1{,}1166.77) = 7.84, p = 0.005$); main effect of track $F(2{,}1213.32) = 4.510, p = 0.011$; reward x track interaction $F(2{,}482.03) = 4.36, p = 0.013$; T1<T2 p = 0.006, T2(HIGH>LOW) p < 0.001). **C**: mean sub-threshold speed (v < 2cm.s$^{-1}$) per lap main effect of reward $F(1{,}1199.13) = 61.01, p < 0.001$; all within track contrasts HIGH<LOW p ≤ 0.002. **D**: time spent immobile anywhere on the track, in seconds, per lap (main effect of reward $F(1{,}1214.42) = 5.39, p = 0.02$; reward x track interaction $F(12{,}732.37) = 13.85, p < 0.001$; T2(HIGH>LOW) p =0.007, T3(HIGH<LOW) p <0.001). **E**: time spent immobile away from reward sites, in seconds, per minute on the track (main effect of reward $F(1{,}67.56) = 8.64, p = 0.004$); T3(HIGH<LOW) p = 0.006). **F**: mean running speed (v > 2cm.s$^{-1}$) per lap (main effect of reward $F(1{,}1225.26) = 28.38, p < 0.001$; main effect of track $F(2{,}1206.23) = 3.141, p =$

$0.04$;reward x track interaction $F(2{,}990.01) = 14.27, p < 0.001$; T3(HIGH>LOW) p <0.001).
**G**: mean + sem speed distributions, in cm.s$^{-1}$, over cumulative stopping time, in seconds, split for LOW and HIGH reward sites.

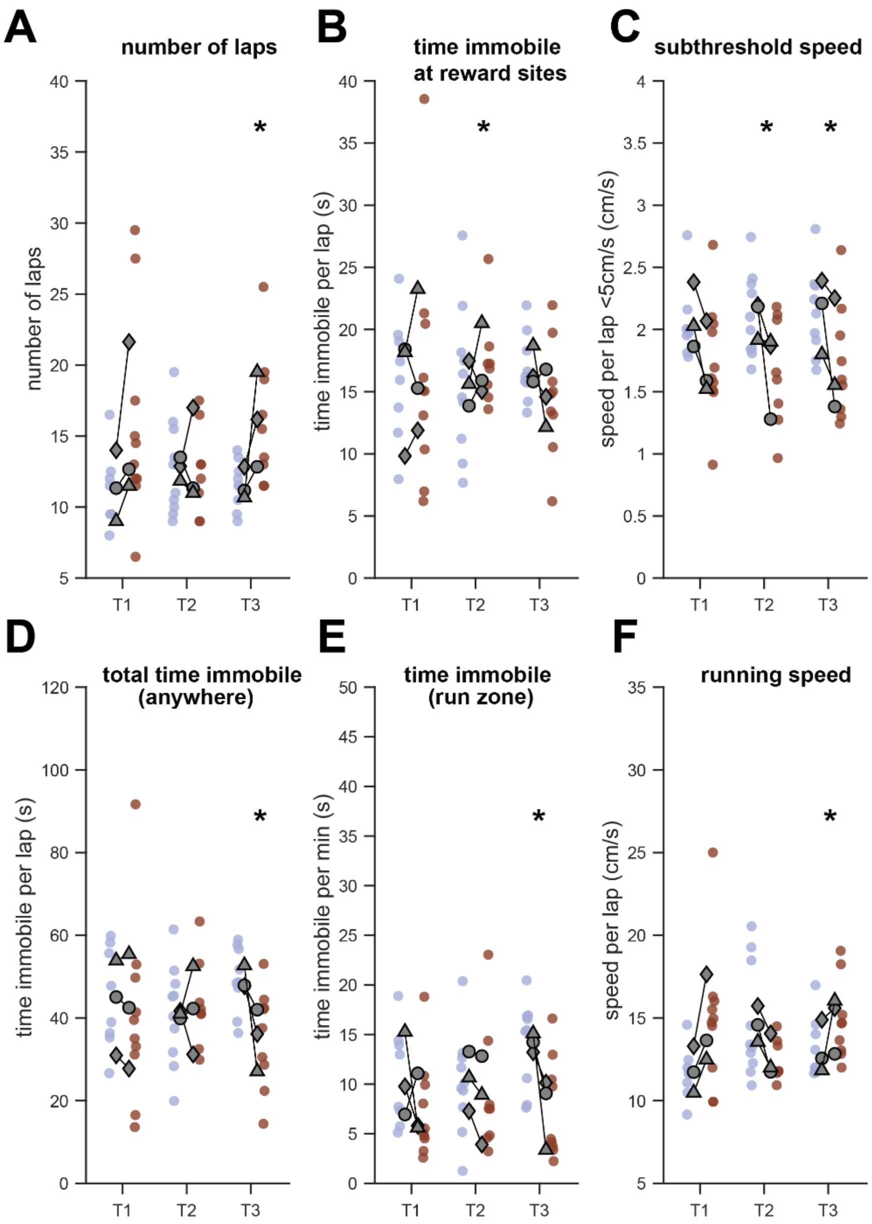

**Fig. S3. Behavioral measures for rats retained for replay analyses. A-F**: Inside markers: rat averages for each measure, split for reward × track identity. Outside round markers are individual datapoints. In brown, HIGH value tracks, in blue LOW value tracks. Rats used for replay analyses have filled in grey symbols. Denoted by (*) significance of simple effects from GLMM ~ reward × track + (1 | rat/session), see corresponding **Table S3 and supplementary text**. **A**: number of laps on each track (main effect of reward $F(1{,}36.11) = 8.502, p = 0.006$; T3(HIGH>LOW) p= 0.012) **B**: time spent immobile at each reward site, in seconds, per lap (main effect of reward $F(1{,}695.5) = 4.615, p = 0.032$; T2(HIGH>LOW) p= 0.006). **C**: mean sub-threshold speed (v < 5 cm.s$^{-1}$) per lap (main effect of reward $F(1{,}700.99) = 38.04, p < 0.001$; trend for T1(HIGH<LOW; T2(HIGH<LOW) p <0.001, T3(HIGH<LOW) p= 0.011). **D**: time spent immobile anywhere on the track, in seconds, per lap (main effect of reward $F(1{,}702.31) = 7.12, p = 0.008$; reward x track interaction $F(2{,}469.01) = 9.501, p < 0.001$; T3(HIGH<LOW) p < 0.001). **E**: time spent immobile away from reward sites, in seconds, per minute (main effect of reward $F(1,45.98) = 6.88, p = 0.012$; T3(HIGH<LOW) p=0.011). **F**: mean running speed ($v > 5 cm.s^{-1}$) per lap (main effect of reward $F(1,694.97) = 13.15, p < 0.001$; reward x track interaction $F(2,694.70) = 15.43, p < 0.001$; T3(HIGH>LOW) p <0.001).

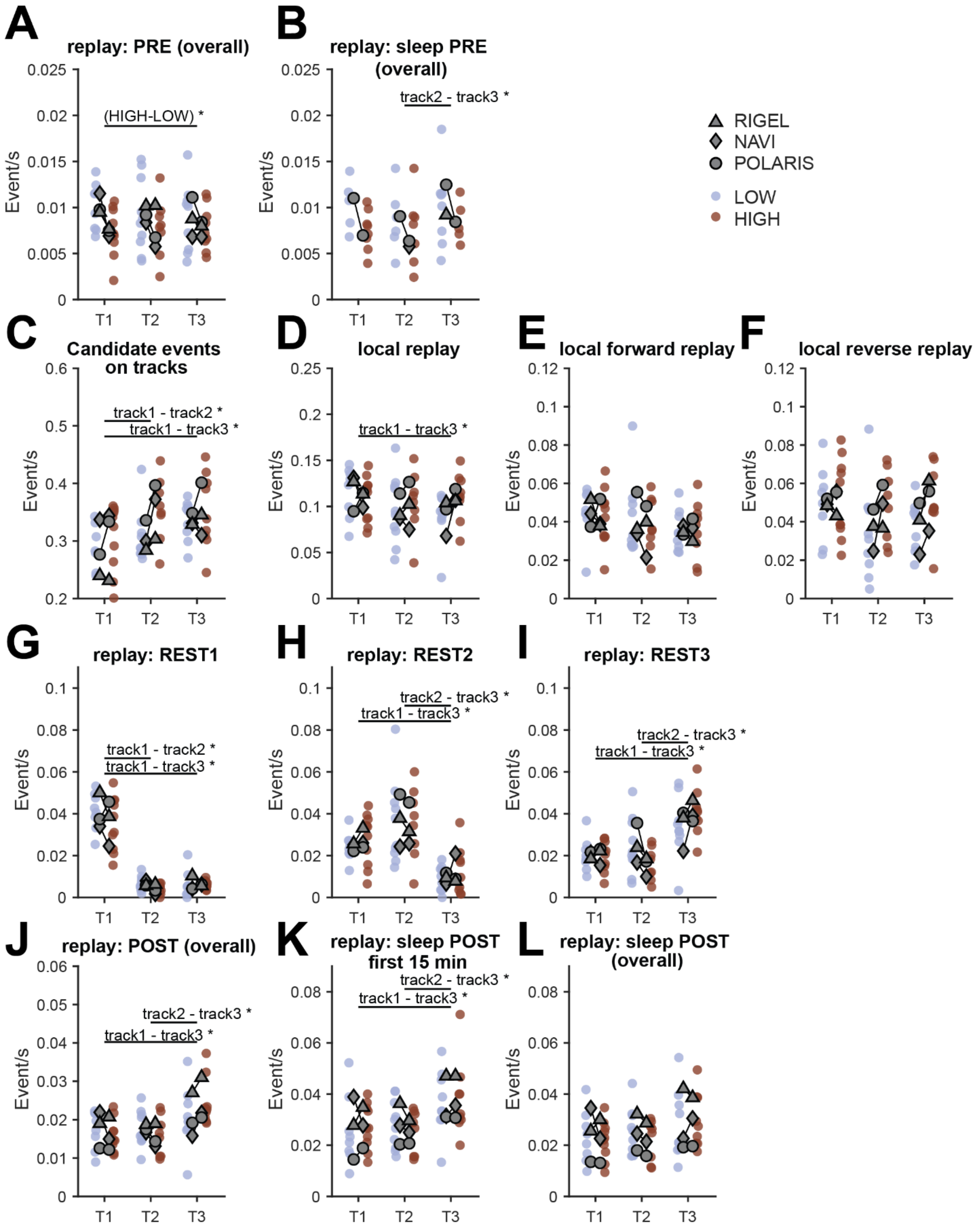
A
replay: PRE (overall)
(HIGH-LOW) *
B
replay: sleep PRE (overall)
track2 - track3 *
RIGEL
NAVI
POLARIS
LOW
HIGH
C
Candidate events on tracks
track1 - track2 *
track1 - track3 *
D
local replay
track1 - track3 *
E
local forward replay
F
local reverse replay
G
replay: REST1
track1 - track2 *
track1 - track3 *
H
replay: REST2
track2 - track3 *
track1 - track3 *
I
replay: REST3
track2 - track3 *
track1 - track3 *
J
replay: POST (overall)
track2 - track3 *
track1 - track3 *
K
replay: sleep POST first 15 min
track2 - track3 *
track1 - track3 *
L
replay: sleep POST (overall)
Event/s
T1
T2
T3

**Fig. S4. Event rates across conditions and epochs.**

Inside markers: rat averages for each measure, split for reward × track identity. Outside round markers are individual datapoints. In brown, HIGH value tracks, in blue LOW value tracks. Denoted by (*) significance of simple effects from GLMM ~ speed + lap number + reward × track + (1 | rat/session), see corresponding **Table S4 and supplementary text**. **A**: offline replay rates during the PRE epoch (main effect of reward $F(1,34.36) = 6.67, p = 0.014$; that did not survive within track simple effects comparisons). **B**: offline replay rates during sleep in PRE. (main effect of track $F(2,26.85) = 3.68, p = 0.038$; T2-T3 p= 0.011). **C**: Online candidate replay rates during episodes (main effect of track $F(2,693.61) = 10.30, p < 0.001$; T1-T2 p= 0.001; T1-T3 p <0.001). **D**: local replay rates during episodes (main effect of track $F(2,691.59) = 3.79, p = 0.023$; reward x track interaction $F(2,354.61) = 3.11, p = 0.046$; T1-T3 p= 0.014; no other contrasts survived correction). **E**: local forward replay rates during episodes (no main effect or interaction). **F**: local reverse replay rates during episodes (no main effect or interaction). **G**: offline replay rates during Rest1 (main effect of track $F(2,45.99) = 130.68, p < 0.001$; T1>T2 p<0.001; T1>T3 p<0.001). **H**: offline replay rates during Rest2 (main effect of track $F(2,31.80) = 19.03, p < 0.001$; T1>T3 p<0.001; T2>T3 p=0.001). **I**: offline replay rates during Rest3 (main effect of track $F(2,29.15) = 21.38, p < 0.001$; T1<T3 p<0.001; T2<T3 p<0.001). **J**: offline replay rate during the POST epoch (main effect of track $F(2,46.003) = 8.87, p = 0.001$; T1<T3 p=0.001; T2<T3 p=0.001). **K**: offline replay rates over the first 15 minutes of sleep during POST (main effect of track $F(2,31.24) = 7.59,\ p = 0.002$; T1<T3 p=0.002; T2<T3 p=0.002). **L**: offline replay rates during sleep in POST (main effect of track $F(2,46) = 3.74, p = 0.031$; trend for T3>T1).

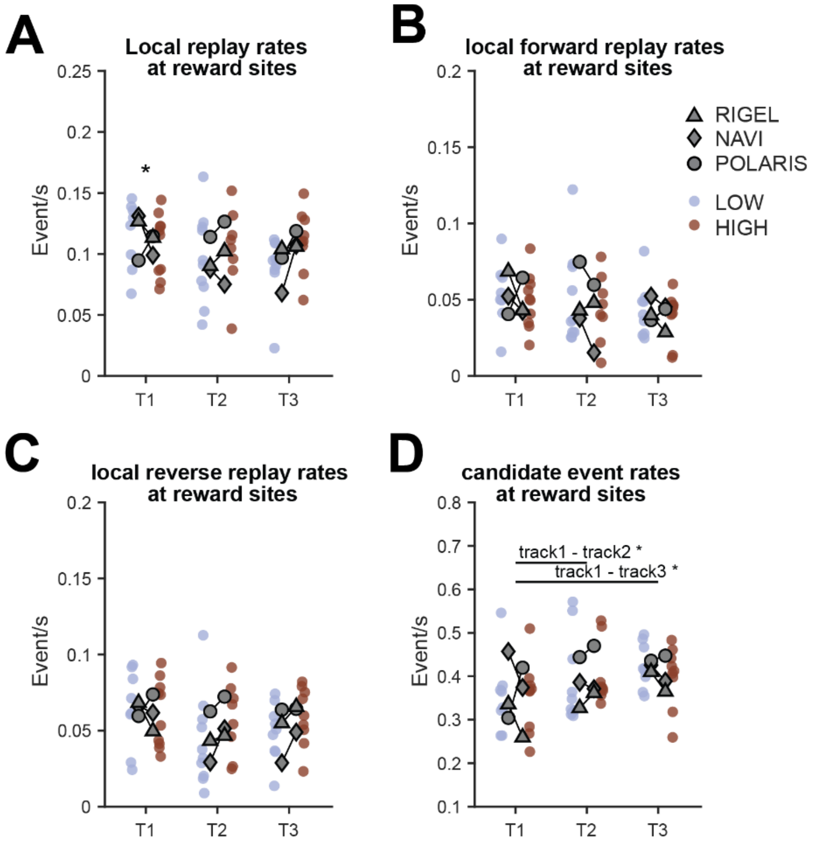

**Fig. S5. Event rates across conditions at reward sites**

Inside markers: rat averages for each measure, split for reward × track identity. Outside round markers are individual datapoints. In brown, HIGH value tracks, in blue LOW value tracks. Rats used for replay analyses have filled in grey symbols. Denoted by (*) significance of simple effects from GLMM ~ speed + lap number + reward × track + (1 | rat/session), **see corresponding Table S5**. **A**: local replay rates at reward sites (main effect of reward $F(1,701.77) = 4.207, p = 0.041$; T1(HIGH<LOW) β=-0.074 p<0.001). **B**: local forward replay rates at reward sites (no main effect of reward, track or their interaction). **C**: local reverse replay rates at reward sites (main effect of track $F(2,692.25) = 3.09, p = 0.046$, reward x track interaction $F(2,249.38)$=3.22, p=0.041; no surviving contrasts, trend T1>T2). D: candidate replay event rates at reward sites (main effect of reward $F(1,697.16) = 5.57, p = 0.018$, main effect of track $F(2,692.20) = 6.056, p = 0.002$; T1<T2 p=0.012, T1<T3 p=0.001; no surviving within track reward contrasts). All replay rates were significantly modulated by subthreshold speed and lap number, see Table S5.

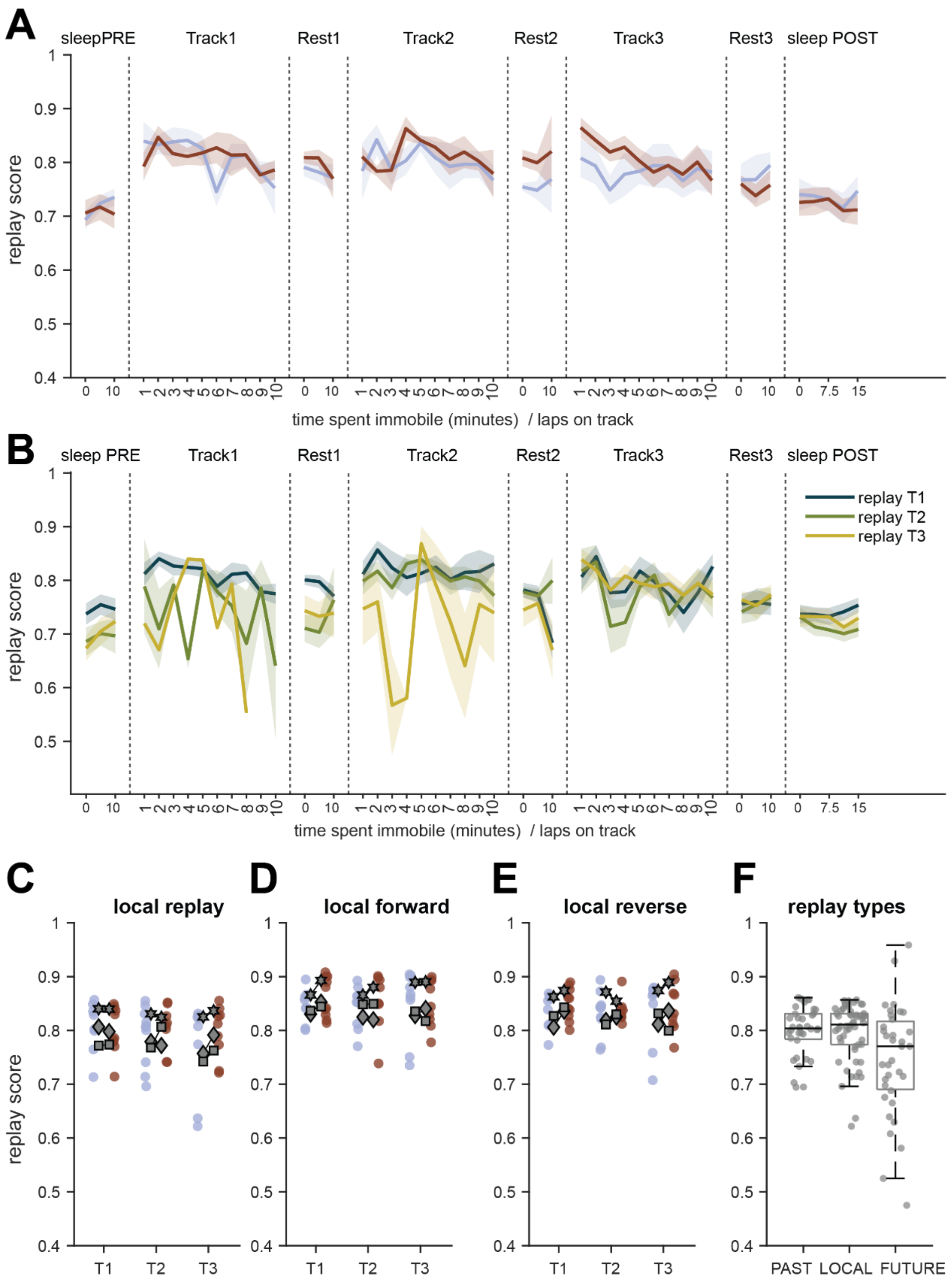

**Fig. S6. Replay quality across conditions and session evolution**

**A,B**: mean and sem of replay quality, as measured by the weighted correlation of the posterior distribution, over time spent immobile (sleep during PRE, Rest1-3, sleep during POST), and over the first 10 laps (Track1-3). **A**: split by reward value. For visualization the replay quality of only one episode (Track 1, 2 or 3) is shown by epoch: the average of all three tracks during sleep

PRE, current track (local replay) for Track1-3 epochs, Track1 during Rest1, Track2 during Rest2, and Track3 during Rest3 and sleep POST. In brown HIGH value tracks, in blue, LOW value tracks. **B**: split by track identity (content of replay). **C-E**: Inside markers: rat averages for each measure, split for reward × track identity. Outside round markers are individual datapoints. In brown, HIGH value tracks, in blue LOW value tracks. Rats used for replay analyses have filled in grey symbols. GLMM $\sim$ reward × track + (1 | rat/session), see corresponding **Table S6**, no main effect of reward, track or their interaction could be observed for panels C-E. **C**: local replay quality. **D**: local forward replay quality. **E**: local reverse replay quality. **F**: boxplots (median + CI) of replay quality for online remote (any), local, and future (any) replay events. Individual points are episodes. Note how the replay quality of future remote replay is lower than past remote and local replay (Kruskal-wallis test,$\chi^2(2) = 9.76, p = 0.007$), Post-hoc multiple comparisons, Tukey corrected (MATLAB's *multcompare*), revealed that both past replay and local replay exhibited higher replay quality than future replay (past vs future: $p = 0.0268,$ local vs future: $p = 0.0105$ whereas replay quality did not differ between past and local replay $p = 0.9966$).

**Table S1-S9.**
See document **StatsSummaryTables.xlsx**
